# Pathway elucidation and microbial synthesis of proaporphine and bis-benzylisoquinoline alkaloids from sacred lotus (*Nelumbo nucifera*)

**DOI:** 10.1101/2022.09.16.508242

**Authors:** Michael E. Pyne, Nicholas D. Gold, Vincent J. J. Martin

**Author notes:** Correspondence and requests for materials should be addressed to: Michael E. Pyne; and Vincent J.J. Martin. Department of Biology, Western University, London, Ontario, Canada (Michael E. Pyne).

## Abstract

Sacred lotus (*Nelumbo nucifera*) has been utilized as a food, medicine, and spiritual symbol for nearly 3,000 years. The medicinal properties of lotus are largely attributed to its unique profile of benzylisoquinoline alkaloids (BIAs), which includes potential anti-cancer, anti-malarial and anti-arrhythmic compounds. BIA biosynthesis in sacred lotus differs markedly from that of opium poppy and other members of the Ranunculales, most notably in an abundance of BIAs possessing the (*R*)-stereochemical configuration and the absence of reticuline, a major branchpoint intermediate in most BIA producers. Owing to these unique metabolic features and the pharmacological potential of lotus, we set out to elucidate the BIA biosynthesis network in *N. nucifera*. Here we show that lotus CYP80G (*Nn*CYP80G) and a superior ortholog from Peruvian nutmeg (*Laurelia sempervirens*; *Ls*CYP80G) stereospecifically convert (*R*)-*N*-methylcoclaurine to the proaporphine alkaloid glaziovine, which is subsequently methylated to pronuciferine, the presumed precursor to nuciferine. While sacred lotus employs a dedicated (*R*)-route to aporphine alkaloids from (*R*)-norcoclaurine, we implemented an artificial stereochemical inversion approach to flip the stereochemistry of the core BIA pathway. Exploiting the unique substrate specificity of dehydroreticuline synthase from common poppy (*Papaver rhoeas*) and pairing it with dehydroreticuline reductase enabled *de novo* synthesis of (*R*)-*N*-methylcoclaurine from (*S*)- norcoclaurine and its subsequent conversion to pronuciferine. We leveraged our stereochemical inversion approach to also elucidate the role of *Nn*CYP80A in sacred lotus metabolism, which we show catalyzes the stereospecific formation of the bis-BIA nelumboferine. Screening our collection of 66 plant *O*-methyltransferases enabled conversion of nelumboferine to liensinine, a potential anti-cancer bis-BIA from sacred lotus. Our work highlights the unique benzylisoquinoline metabolism of *N. nucifera* and enables the targeted overproduction of potential lotus pharmaceuticals using engineered microbial systems.

## INTRODUCTION

As the national flower of both India and Vietnam, sacred lotus (*Nelumbo nucifera*) has played a major historical role in Asian food, medicine, culture, and religion. Seeds of *N. nucifera* have been consumed for approximately 3,000 years, while most parts of the plant have been used in traditional Chinese medicine to treat fever, diarrhea, cholera, inflammation, and other ailments. The potential health benefits of lotus are largely ascribed to the plant’s unique collection of benzylisoquinoline alkaloids (BIAs), which includes many 1-benzylisoquinolines, 15 aporphine alkaloids, seven bis-BIAs, and one unusual tri-BIA (Menéndez-Perdomo and Facchini, 2018; Yang et al., 2018). BIAs produced by sacred lotus exhibit a range of promising pharmacological properties for potential treatment of cancer (neferine and liensinine), malaria (roemerine), arrhythmia (neferine), HIV (norcoclaurine), obesity (pronuciferine), and fungal infection (roemerine). A recent screen of nearly 200 natural products identified the bis-BIA neferine as a pan-coronavirus entry inhibitor with promising *in vitro* activity against the SARS-CoV-2 virus (He et al., 2021).

The sacred lotus BIA metabolic network and its associated metabolites exhibit a number of structural and stereochemical features that are not found in members of the Ranunculales. (*S*)- Reticuline is the major branchpoint intermediate in most BIA-producing plants, giving rise to more than 2,000 derivatized structures within the morphinan, aporphine, protoberberine, phthalideisoquinoline, and benzophenanthridine downstream branches. However, reticuline has not been isolated from sacred lotus, implying that aporphine and bis-BIA pathways branch from *N*-methylcoclaurine or armepavine in *N. nucifera*. Additionally, aporphine alkaloids from *N. nucifera* lack 4ʹ-substitution of the benzyl moiety that derives from 4-hydroxyphenylacetaldehyde in nearly all BIAs elucidated to date (Pyne et al., 2020). An exception is pronuciferine, an unusual proaporphine alkaloid synthesized by lotus, which possesses a 4ʹ-substituted benzyl group, implying the presence of a novel uncharacterized dehydration reaction involved in the conversion of pronuciferine to nuciferine (Menéndez-Perdomo and Facchini, 2018).

Beyond structural features, BIAs synthesized by sacred lotus exhibit unique stereochemistry compared to other BIA-producing plants. Most BIAs found in nature are (*S*)- conformers owing to the stereospecific synthesis of (*S*)-norcoclaurine by all norcoclaurine synthase (NCS) orthologs characterized to date. In contrast, sacred lotus synthesizes both (*R*)- and (*S*)-conformers of norcoclaurine. Although the *N. nucifera* genome contains at least five NCS isoforms, a dedicated (*R*)- or (*R*,*S*)-yielding NCS has yet to be identified (Menéndez-Perdomo and Facchini, 2018). (*R*)-*N*-methylcoclaurine and both (*R*)- and (*S*)-stereoisomers of armepavine have also been isolated from sacred lotus (Do et al., 2013; Ka et al., 2010; Kashiwada et al., 2005; Nishibe et al., 1986), suggesting a lack of stereospecificity of BIA methyltransferases from *N. nucifera*. Two such lotus methyltransferases involved in the conversion of norcoclaurine to coclaurine (*Nn*6OMT) and *N*-methylcoclaurine to armepavine (*Nn*OMT5) were recently characterized and shown to methylate both (*S*)- and (*R*)-substrates (Menéndez-Perdomo and Facchini, 2020). All aporphine alkaloids isolated from *N. nucifera* to date possess the (*R*)-configuration, while bis-BIAs from lotus arise from both (*R*)- and (*S*)- precursors or two (*R*)-monomers (Menéndez-Perdomo and Facchini, 2018), highlighting a distinct preference for (*R*)-conformers in downstream branches of the lotus BIA network. These unique structural and stereochemical features of BIA metabolites from *N. nucifera* provide substantial rationale for elucidating BIA biosynthesis in sacred lotus and engineering its production in yeast.

Previously, we constructed a yeast BIA platform with capacity to synthesize more than 4.5 g/L of plant BIAs (Pyne et al., 2020). Owing to the pharmaceutical potential and unique features of BIA metabolism in sacred lotus, we wished to leverage our BIA production platform for the discovery and engineering of lotus BIA biosynthesis in yeast. In this work we design and implement an artificial pathway to invert the stereochemistry of our high titer BIA production platform. Flipping the stereochemistry of the core BIA pathway enabled access to the bis-BIA and aporphine branches of sacred lotus secondary metabolism. We show that *Nn*CYP80G and *Nn*CYP80A (also referred to as *Nn*CYP80Q1 and *Nn*CYP80Q2) (Nelson and Schuler, 2013) are stereospecific for (*R*)-substrates and catalyze committed steps in the aporphine and bis-BIA branches in sacred lotus. Our work sheds light on the unique BIA metabolism of *N. nucifera* and will enable the pharmacological exploration and overproduction of untapped BIA structures synthesized by sacred lotus.

## RESULTS

### *De novo* production of armepavine

To begin reconstructing the lotus aporphine biosynthesis pathway in yeast, we selected the (*S*)- norcoclaurine-producing strain LP603 as the basis for our work. This strain harbors an overexpressed and deregulated precursor pathway (*ARO4^FBR^*, *ARO2*, *ARO7^FBR^*, *TYR1*, *ARO10*), two copies of an optimized (*S*)-norcoclaurine synthase gene (*CjNCSΔN*), and deletion of several key host genes encoding oxidoreductases (*ari1*Δ *adh6*Δ *ypr1*Δ *ydr541c*Δ *gre2*Δ). LP478, the parent of LP603, produces more than 1 g/L of (*S*)-norcoclaurine in a sucrose pulsed fed-batch fermentation (Pyne et al., 2020).

Armepavine, a key BIA intermediate in sacred lotus, is synthesized from norcoclaurine via successive action of 6-*O*-methyltransferase (6OMT), coclaurine-*N*-methyltransferase (CNMT), and 7-*O*-methyltransferase (7OMT). Recently, OMT5 from *N. nucifera* (*Nn*OMT5) was characterized from sacred lotus and shown to exhibit 7OMT activity on *N*-methylcoclaurine, yielding armepavine (Menéndez-Perdomo and Facchini, 2020). To engineer LP603 for high-level armepavine production, we introduced a chimeric armepavine production pathway consisting of 6OMT and CNMT from *Papaver somniferum* (*Ps*6OMT and *Ps*CNMT, respectively), in addition to *Nn*OMT5 (**Fig. 1a**). 6OMT and CNMT variants from opium poppy (*P. somniferum*) dramatically outperformed orthologous isoforms from *N. nucifera* (*Nn*6OMT1- 4, *Nn*CNMT1, and *Nn*CNMT3) (**Supplementary Fig. 1**). Strain LP609 synthesized 152 mg/L armepavine in small-scale microtiter plate cultivations, corresponding to a 72% consumption of (*S*)-norcoclaurine (**9**) (**Fig. 1b**). A minor amount of (*S*)-coclaurine (**10**) was produced by strain LP609, while a substantial proportion of (*S*)-*N*-methylcoclaurine (**11**) accumulated, indicating incomplete conversion to armepavine by *Nn*OMT5.

**Figure 1.**
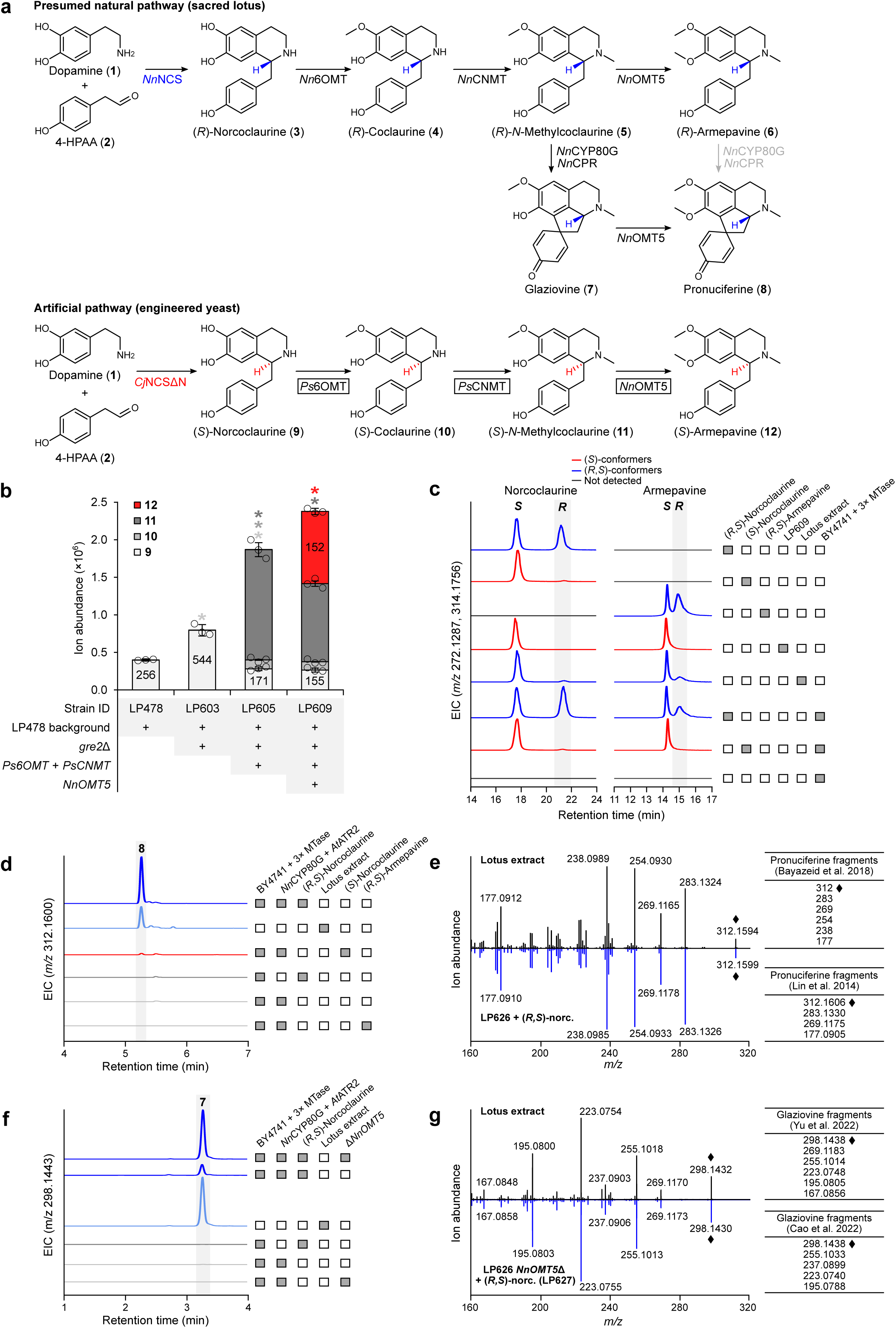
*Nn*CYP80G is a stereospecific proaporphine synthase. **a,** Proposed natural and artificial routes to armepavine, glaziovine, and pronuciferine in sacred lotus and engineered yeast, respectively. Lotus possesses five NCS isoforms for synthesis of (*R*)-norcoclaurine, which is subsequently methylated to (*R*)-armepavine. Yeast strains utilized in this study employ a truncated NCS variant from *C. japonica* (*Cj*NCSΔN), which yields exclusively (*S*)- norcoclaurine. *Ps*6OMT, *Ps*CNMT, and *Nn*OMT5 (3× MTase) involved in the conversion of (*S*)- norcoclaurine to (*S*)-armepavine are boxed. **b,** Analysis of BIA production by engineered strains. Introduction of *Nn*OMT5 to strain LP603 enables high titer production of armepavine. Peak area is shown for all pathway intermediates and absolute titers are depicted in mg/L for (*S*)- norcoclaurine and (*S*)-armepavine. Asterisk (*) denotes a significant increase or decrease (*P* < 0.05) in ion abundance relative to the parent strain. Error bars represent the mean ± s.d. of *n* = 3 independent biological samples. Statistical differences between control and derivative strains were tested using two-tailed Student’s *t*-test. **c,** Chiral analysis of norcoclaurine and armepavine synthesized *de novo* or from norcoclaurine-supplemented strains of yeast. Yeast-derived norcoclaurine was compared against authentic (*S*)-norcoclaurine standard and (*R*,*S*)- norcoclaurine generated by the phosphate-catalyzed condensation of dopamine and 4-HPAA. Commercial (*S*)-norcoclaurine standard contains a small proportion of (*R*)-norcoclaurine, as described previously (Pyne et al., 2020). Yeast-derived armepavine was compared against authentic (*R*,*S*)-armepavine standard, sacred lotus extract, and (*S*)-armepavine generated from feeding (*S*)-norcoclaurine to a BY4741 strain harboring *Ps*6OMT, *Ps*CNMT, and *Nn*OMT5 (3× MTase). (*R*)-Conformers of norcoclaurine and armepavine are shaded. **d,** Ion-extracted LC–MS chromatograms of engineered yeast strains harboring a partial pronuciferine pathway (*Ps*6OMT-*Ps*CNMT-*Nn*OMT5-*Nn*CYP80G-*At*ATR2; LP626) fed with (*R*,*S*)-norcoclaurine, (*S*)- norcoclaurine, or (*R*,*S*)-armepavine. LP626 is unable to synthesize BIAs *de novo* due to lack of an NCS enzyme and a dopamine biosynthesis pathway. Retention time of yeast-derived pronuciferine (dark blue) was compared against sacred lotus extract (light blue). **e,** MS/MS fragmentation of pronuciferine from sacred lotus extract and strain LP626 fed with (*R,S*)- norcoclaurine. Pronuciferine fragmentation is compared against two prior reports using authentic pronuciferine standard (Bayazeid et al., 2018; Lin et al., 2014). **f,** Ion-extracted LC-MS chromatograms of strain LP626 and its *NnOMT5*Δ derivative (LP627) fed with (*R*,*S*)- norcoclaurine. Retention time of yeast-derived glaziovine (dark blue) was compared against sacred lotus extract (light blue). **g,** MS/MS fragmentation of glaziovine from sacred lotus extract and strain LP627 fed with (*R,S*)-norcoclaurine. Glaziovine fragmentation is compared against two prior reports using authentic glaziovine standard (Cao et al., 2022; Yu et al., 2022).

### *Nelumbo nucifera* CYP80G is a stereospecific proaporphine synthase

Having engineered *de novo* synthesis of armepavine (LP609), we set out to characterize the predicted *N. nucifera* aporphine synthase (*Nn*CYP80G) in yeast. However, introduction of *Nn*CYP80G and a cytochrome P450 reductase (CPR) from *Arabidopsis thaliana* (*At*ATR2; strain LP624) failed to alter the metabolite profile of LP609. Swapping *At*ATR2 for a CPR from *Hypericum calycinum* (*Hc*CPR) or including a cytochrome *b5* variant from *Artemisia annua* (*Aa*CYB5) also failed to yield new metabolite peaks consistent with proaporphine or aporphine metabolites.

Sacred lotus synthesizes both (*S*)- and (*R*)-conformers of norcoclaurine and armepavine (Menéndez-Perdomo and Facchini, 2018), leading us to speculate that *Nn*CYP80G might be stereospecific. Our armepavine production strain (LP609) employs *Cj*NCSΔN from *Coptis japonica*, which we showed yields exclusively (*S*)-norcoclaurine (Pyne et al., 2020), implying that armepavine produced by LP609 is the (*S*)-conformer. To clarify the stereochemistry of BIAs synthesized by strain LP609, we performed chiral analysis on norcoclaurine and armepavine, which were found to be exclusively (*S*)-conformers (**Fig. 1c**). We also detected both (*S*)- and (*R*)- stereoisomers of norcoclaurine and armepavine from commercial extracts of lotus, which is consistent with previous metabolite characterization studies (Menéndez-Perdomo and Facchini, 2018).

Because LP609 synthesizes exclusively (*S*)-BIAs, yet *N. nucifera* produces both (*S*)- and (*R*)-BIAs, we speculated that *Nn*CYP80G is stereospecific for (*R*)-substrates. To establish the stereospecificity of *Nn*CYP80G, we introduced a partial aporphine production pathway (*Ps*6OMT-*Ps*CNMT-*Nn*OMT5-*Nn*CYP80G-*At*ATR2) to an unmodified BY4741 laboratory strain lacking an NCS enzyme and a dopamine biosynthesis pathway (yielding strain LP626). LP626 is unable to synthesize BIAs *de novo*, thus enabling its use as a host for feeding various stereoisomers of norcoclaurine. Feeding strain LP626 with (*R*,*S*)-norcoclaurine (*rac*- norcoclaurine), generated by the *in vitro* phosphate-catalyzed condensation of dopamine (**1**) and 4-hydroxyphenylacetaldehyde (**2**) (Pesnot et al., 2012), yielded (*R*)-armepavine, in addition to (*S*)-armepavine, demonstrating three-step methylation of (*R*)-norcoclaurine (**3**) to (*R*)- armepavine (**6**) by *Ps*6OMT, *Ps*CNMT, and *Nn*OMT5 (**Fig. 1c**). Importantly, feeding *rac*- norcoclaurine to strain LP626 resulted in the formation of a new LC-MS metabolite peak consistent with pronuciferine (**8**) ([M + H]^+^ = 312.1600 Da) (**Fig. 1d**; dark blue trace). Omitting *Nn*CYP80G failed to generate the tentative pronuciferine LC-MS product. Unexpectedly, feeding (*S*)-norcoclaurine standard to strain LP626 also generated pronuciferine, yet with a peak area approximately 98% smaller than the corresponding peak derived from supplemented *rac*- norcoclaurine (**Fig. 1d**; red trace). Chiral analysis of commercial (*S*)-norcoclaurine revealed that the standard was contaminated with a small quantity of (*R*)-norcoclaurine (**Fig. 1c**), as also shown in a prior study (Pyne et al., 2020), providing explanation for the low level of pronuciferine synthesized by LP626 from supplemented (*S*)-norcoclaurine. Supplementing phosphate buffer in place of *rac*-norcoclaurine and (*S*)-norcoclaurine failed to generate the tentative pronuciferine LC-MS product (**Fig. 1d**).

Although an authentic standard of pronuciferine is not available, we detected the same tentative LC-MS peak from commercial lotus leaf extract. Pronuciferine from lotus extract exhibited the same retention time and exact mass as the tentative peak derived from supernatants of strain LP626 supplemented with *rac*-norcoclaurine (**Fig. 1d**; light blue trace). Targeted fragmentation of the tentative pronuciferine peak from lotus extract and supernatants of LP626 fed with *rac*-norcoclaurine yielded identical fragmentation spectra (**Fig. 1e**), which were in agreement with published spectra derived from authentic pronuciferine (Bayazeid et al., 2018; Lin et al., 2014).

Based on our *rac*-norcoclaurine feeding results (**Fig. 1d**) and prior studies detailing alkaloid isolation (Bishayee et al., 2022; Cao et al., 2022; Yu et al., 2022) and *in vivo* tracer experiments using sacred lotus (Barton et al., 1967; Haynes et al., 1965), we deduced two theoretical routes to pronuciferine (**Fig. 1a**). Because pronuciferine exhibits the same methylation pattern as armepavine, one route involves the direct conversion of (*R*)-armepavine to pronuciferine by *Nn*CYP80G and ATR2. However, based on the mechanism of proaporphine synthesis (Barton and Cohen, 1957), phenol coupling is unable to proceed in 7-*O*-methylated substrates. This mechanism was validated using *in vivo* tracer studies (Barton et al., 1967; Haynes et al., 1965), in which labelled 7-*O*-methylated substrates, such as isococlaurine and armepavine, were not converted to corresponding proaporphine products, while labelled norcoclaurine, coclaurine and *N*-methylcoclaurine substrates were efficiently incorporated.

An alternative route to pronuciferine involves conversion of (*R*)-*N*-methylcoclaurine to the proaporphine glaziovine (**7**) and subsequent methylation by *Nn*OMT5, yielding pronuciferine. To clarify the preferred substrate of *Nn*CYP80G and the major route to pronuciferine in sacred lotus, we supplemented (*R*,*S*)-armepavine to strain LP626 harboring a partial proaporphine production pathway (*Ps*6OMT-*Ps*CNMT-*Nn*OMT5-*Nn*CYP80G-*At*ATR2). Whereas feeding 2× SC medium containing (*R*,*S*)-norcoclaurine to LP626 facilitated synthesis of pronuciferine, supplementing (*R*,*S*)-armepavine failed to yield pronuciferine (**Fig. 1d**), corroborating prior evidence that armepavine is not a precursor to proaporphines (Barton et al., 1967). We were also unable to detect synthesis of pronuciferine from supplemented (*R*,*S*)- armepavine by switching to YPD medium or by increasing the pH of 2× SC or YPD medium to 7.0, as increasing culture pH improves uptake of BIA metabolites by *S. cerevisiae* (Fossati et al., 2015; Pyne et al., 2016).

Owing to the inability of *Nn*CYP80G to convert exogenous (*R*,*S*)-armepavine to pronuciferine, we investigated *Nn*CYP80G activity on *N*-methylcoclaurine, which lacks 7-*O*- methylation, by first deleting the gene encoding *NnOMT5* in LP626, yielding strain LP627. In the absence of *Nn*OMT5, LP627 accumulates (*R*,*S*)-*N*-methylcoclaurine from supplemented (*R*,*S*)-norcoclaurine and is thus unable to synthesize pronuciferine. Feeding (*R*,*S*)-norcoclaurine to LP627 resulted in the formation of a new LC-MS metabolite peak consistent with glaziovine (**7**) ([M + H]^+^ = 298.1443 Da) (**Fig. 1d**; dark blue trace), a propaporphine alkaloid from *N. nucifera* (Cao et al., 2022; Yu et al., 2022). Omitting *Nn*CYP80G or (*R*,*S*)-norcoclaurine failed to generate the tentative glaziovine LC-MS product. We also detected an LC-MS peak from commercial lotus leaf extract with the same retention time and exact mass as the tentative glaziovine peak derived from supernatants of strain LP627 supplemented with *rac*-norcoclaurine (**Fig. 1f**; light blue trace). Targeted fragmentation of the tentative glaziovine peak from supernatants of LP627 fed with *rac*-norcoclaurine yielded fragmentation spectra that were in agreement with published spectra derived from authentic glaziovine (Cao et al., 2022; Yu et al., 2022) (**Fig. 1g**). Importantly, glaziovine accumulation decreased markedly in the presence of *Nn*OMT5, which coincided with the formation of pronuciferine, confirming that proaporphine formation precedes 7-*O*-methylation. Collectively, these data demonstrate that *Nn*CYP80G stereospecifically converts (*R*)-*N*-methylcoclaurine to glaziovine in the committed step of the sacred lotus aporphine biosynthesis pathway.

Based on BIA alkaloids isolated from *N. nucifera*, synthesis of nuciferine arises from an unknown dehydration reaction from pronuciferine (Menéndez-Perdomo and Facchini, 2018). Recently, a study of alkaloids from *N. nucifera* identified a cluster of 90 genes closely associated with alkaloid content, including *Nn*CYP80G and several BIA methyltransferases (Yang et al., 2017). However, most of the 90 identified genes do not encode biosynthetic enzymes or encode biosynthetic enzymes, such as methyltransferases, not involved in the conversion of pronuciferine to nuciferine. In an attempt to identify the missing activity, we selected 10 of these candidate biosynthetic genes from lotus (NNU05340, NNU08935, NNU17596, NNU01031, NNU20422, NNU17669, NNU24387, NNU12203, NNU07402, and NNU20868), which were integrated into a pronuciferine-producing strain (LP634). None of the candidate lotus genes facilitated accumulation of nuciferine or consumption of pronuciferine.

### An artificial *de novo* route to (*R*)-*N*-methylcoclaurine and (*R*)-armepavine using reticuline epimerase

Having established a role for *Nn*CYP80G in proaporphine alkaloid biosynthesis using supplementation assays, we aimed to engineer yeast for *de novo* synthesis of pronuciferine. Because lotus ostensibly synthesizes pronuciferine from (*R*)-norcoclaurine in a dedicated (*R*)- specific pathway (**Fig. 1a**) (Menéndez-Perdomo and Facchini, 2018), reconstructing the natural lotus route in yeast would entail screening NCS orthologs from *N. nucifera* and swapping *Cj*NCSΔN for an (*R*)-specific isoform. However, *Cj*NCSΔN dramatically outperformed all NCS variants screened from a diverse array of plants (Bourgeois et al., 2018; Grewal et al., 2021; Pyne and Martin, 2022), enabling high titers (> 4.5 g/L) of BIAs (Pyne et al., 2020).

We searched for an alternative route to (*R*)-*N*-methylcoclaurine (**5**) and (*R*)-armepavine (**6**) from (*S*)-norcoclaurine (**9**) and shifted our focus to reticuline epimerase (REPI), an enzyme involved in the committed step of the morphinan BIA pathway (Farrow et al., 2015; Galanie et al., 2015; Winzer et al., 2015). REPI catalyzes the two-step stereochemical inversion of (*S*)- to (*R*)-reticuline via the intermediate 1,2-dehydroreticuline. REPI is composed of dehydroreticuline synthase (DRS) and dehydroreticuline reductase (DRR) domains, which catalyze the respective synthesis and reduction of 1,2-dehydroreticuline. While DRS from opium poppy (*P. somniferum*; *Ps*DRS) is specific for (*S*)-reticuline, DRS from common poppy (*P. rhoeas*; *Pr*DRS) exhibits unique preference for (*S*)-*N*-methylcoclaurine (Farrow et al., 2015), the precursor to armepavine in sacred lotus. We speculated that screening diverse pairs of DRS and DRR orthologs might identify a synthetic DRS-DRR couple capable of catalyzing the stereochemical inversion of (*S*)- to (*R*)-*N*-methylcoclaurine (**Fig. 2a**). A concurrent study employed a similar approach, albeit using supplementation with L-DOPA (Payne et al., 2021). Instead, we wished to develop a fully *de novo* route to (*R*)-*N*-methylcoclaurine and pronuciferine without supplementation of BIA pathway precursors.

**Figure 2.**
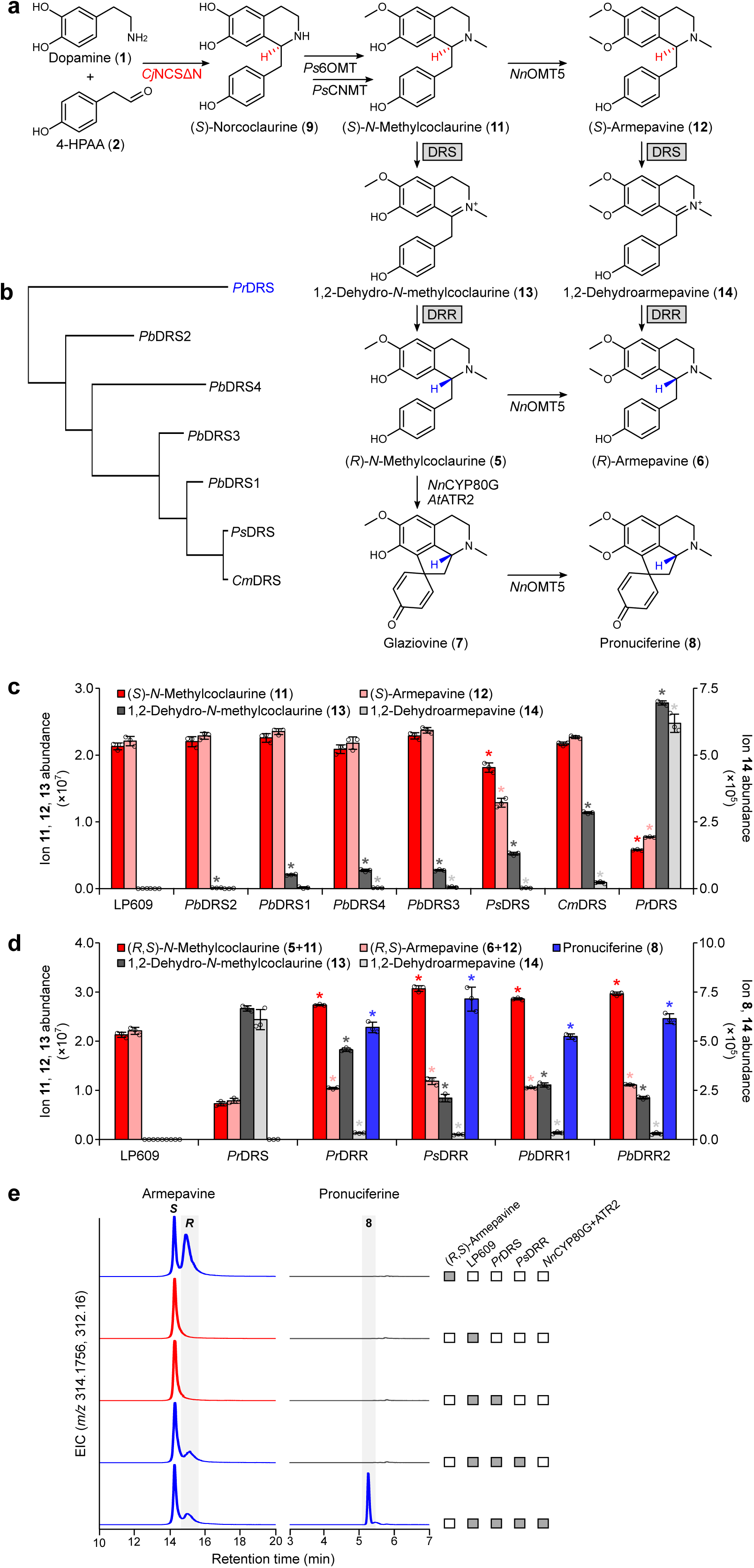
Screening poppy dehydroreticuline synthase (DRS) and dehydroreticuline reductase (DRR) orthologs for *de novo* synthesis of pronuciferine from (*R*)-BIA conformers. **a,** Artificial *de novo* routes to (*R*)-*N*-methylcoclaurine and pronuciferine via stereochemical inversion. (*S*)-Enantiomers are converted to corresponding 1,2-dehydro intermediates using DRS, which are subsequently reduced to (*R*)-conformers using DRR. **b,** Protein phylogenetic tree of DRS orthologs from various poppy species. Abbreviations: *Pr*, *Papaver rhoeas* (common poppy); *Pb*, *Papaver bracteatum* (Iranian poppy); *Ps*, *Papaver somniferum* (opium poppy); *Cm*, *Chelidonium majus* (greater celandine). **c,** Synthesis of 1,2- dehydro-*N*-methylcoclaurine and 1,2-dehydroarmepavine by poppy DRS variants. DRS- encoding genes were integrated into an (*S*)-armepavine-producing strain (LP609) and formation of 1,2-dehydro intermediates was assayed by LC-MS. Asterisk (*) denotes a significant increase or decrease (*P* < 0.05) in ion abundance relative to the precursor strain (LP609). **d,** Reduction of 1,2-dehydro-*N*-methylcoclaurine and 1,2-dehydroarmepavine by poppy DRR orthologs. Plasmid-encoded DRR genes were introduced to LP609 harboring *Pr*DRS, *Nn*CYP80G, and *At*ATR2, and consumption of 1,2-dehydro intermediates was assayed by LC-MS. *N*-Methylcoclaurine and armepavine are reported as combined (*R*)- and (*S*)-conformers. Error bars represent the mean ± s.d. of *n* = 3 independent biological samples. Asterisk (*) denotes a significant increase or decrease (*P* < 0.05) in ion abundance relative to the precursor strain harboring *Pr*DRS, *Nn*CYP80G, and *At*ATR2. Statistical differences between control and derivative strains were tested using two-tailed Student’s *t*-test. **e,** Chiral analysis of armepavine synthesized by strains harboring *Pr*DRS or *Pr*DRS+*Ps*DRR. Implementation of *Pr*DRS+*Ps*DRR in conjunction with *Nn*CYP80G+*At*ATR2 enables stereochemical inversion and concurrent synthesis of pronuciferine. Pronuciferine was separated on a standard (non-chiral) C18 column.

To engineer a *de novo* stereochemical inversion path to pronuciferine, we first screened several DRS variants for activity on (*S*)-*N*-methylcoclaurine. This screen included *Pr*DRS, *Ps*DRS, DRS from *Chelidonium majus* (*Cm*DRS) and four isoforms from Iranian poppy (*P. bracteatum*; *Pb*DRS1-4). *Cm*DRS and *Pb*DRS1-4 share 86-99% identity with the canonical *Ps*DRS variant, whereas *Pr*DRS shares only 79% identity to *Ps*DRS (**Fig. 2b**). In agreement with a prior study (Farrow et al., 2015), *Pr*DRS exhibited remarkable activity on (*S*)-*N*- methylcoclaurine, yielding 1,2-dehydro-*N*-methylcoclaurine (**13**) ([M + H]^+^ = 298.1443 Da) (**Fig. 2c**).

In addition to its reported activity on (*S*)-*N*-methylcoclaurine, we screened *Pr*DRS and other DRS variants on (*S*)-armepavine. Remarkably, *Pr*DRS was the only DRS variant that converted (*S*)-armepavine to 1,2-dehydroarmepavine (**14**) ([M + H]^+^ = 312.1600 Da) (**Fig. 2c**). Implementation of *Pr*DRS converted nearly 70% of total (*S*)-substrate [combined (*S*)-*N*- methylcoclaurine and (*S*)-armepavine] to dehydro-intermediates. Assuming equivalent LC-MS response factors, the strain harboring *Pr*DRS produced 45-fold more 1,2-dehydro-*N*- methylcoclaurine than 1,2-dehydro-armepavine, although consumption of (*S*)-*N*- methylcoclaurine reduces the pool of available (*S*)-armepavine. Aside from *Pr*DRS, *Ps*DRS consumed 29% of (*S*)-substrates, while the remaining DRS orthologs converted a very small proportion of available (*S*)-conformers to the corresponding dehydro-derivatives.

Using *Pr*DRS for synthesis of 1,2-dehydro-BIAs, we next screened DRR variants for reduction of 1,2-dehydro-*N*-methylcoclaurine and 1,2-dehydroarmepavine. Our screen included *Pr*DRR, *Ps*DRR, and two isoforms of DRR from *P. bracteatum* (*Pb*DRR1 and *Pb*DRR2) (**Fig. 2d**). Introduction of all four DRR orthologs facilitated the near-complete consumption of 1,2- dehydroarmepavine. While none of the DRR variants completely consumed 1,2-dehydro-*N*- methylcoclaurine, strains harboring *Ps*DRR or *Pb*DRR2 each reduced 68% of available substrate. Because authentic (*R*,*S*)-armepavine is commercially available, we utilized (*R*)- armepavine to approximate (*R*)-*N*-methylcoclaurine synthesis, as *Nn*OMT5 catalyzes the methylation of both (*S*)- and (*R*)-conformers of *N*-methylcoclaurine (**Fig. 1c**). Chiral LC-MS analysis of supernatants derived from strains harboring *Pr*DRS and *Ps*DRR revealed synthesis of (*R*)-armepavine, while strains lacking a DRR domain produced exclusively (*S*)-armepavine (**Fig. 2e**). Importantly, inclusion of *Nn*CYP80G and ATR2 along with *Pr*DRS and *Ps*DRR facilitated *de novo* production of pronuciferine (**Fig. 2d,e**), further establishing the involvement of (*R*)-BIAs in its synthesis. Collectively, these modifications establish an artificial *de novo* route to (*R*)-*N*-methylcoclaurine and pronuciferine via (*S*)-norcoclaurine.

### Enhancing CYP80G-CPR coupling and probing activity of *Nn*CYP80A

After establishing the role of *Nn*CYP80G in proaporphine alkaloid biosynthesis, we attempted to improve its activity through better coupling to a CPR partner. We assembled a combinatorial CYP80-CPR-CYB5 library by pairing different combinations of plant CPRs and CYB5s with *Nn*CYP80G or *Ls*CYP80G, a predicted CYP80G ortholog from *Laurelia sempervirens* identified from our in-house collection of plant CYPs. We also included in this assay *Nn*CYP80A, a lotus enzyme possessing 80% identity to *Nn*CYP80G. Genes encoding *NnCYP80G* and *NnCYP80A* are adjacent within the lotus genome, suggesting tandem duplication and a potential role for *Nn*CYP80A in BIA biosynthesis (Menéndez-Perdomo and Facchini, 2018). Previous studies have implicated *Nn*CYP80A in bis-BIA biosynthesis, yet its activity has not been experimentally validated. Because all bis-BIAs characterized from lotus derive from at least one (*R*)-monomer, our ability to synthesize both (*S*)- and (*R*)-conformers using *Pr*DRS and DRR provides an opportunity to fully explore *Nn*CYP80A activity.

We first assembled a CYP80-CPR-CYB5 library consisting of 24 combinations of one CYP80 (*Nn*CYP80A, *Nn*CYP80G, and *Ls*CYP80G), each paired with one of four CPRs and one of four CYB5s (**Fig. 3a**). We then transformed our 24 strain CYP80-CPR-CYB5 library with a plasmid that expresses both *PrDRS* and *PsDRR* genes for conversion of (*S*)-intermediates to (*R*)- conformers. As we demonstrated above (**Fig. 1d**), all *Nn*CYP80G strains expressing *PrDRS* and *PsDRR* produced pronuciferine (**Fig. 3b**). However, all strains harboring *Ls*CYP80G synthesized more pronuciferine than isogenic strains possessing *Nn*CYP80G. Selection of both CPR and CYB5 variant had a significant effect on pronuciferine production, whereby pronuciferine output was greatest when *Ls*CYP80G was coupled with *Cc*CPR2 and *At*CYB5D.

**Figure 3.**
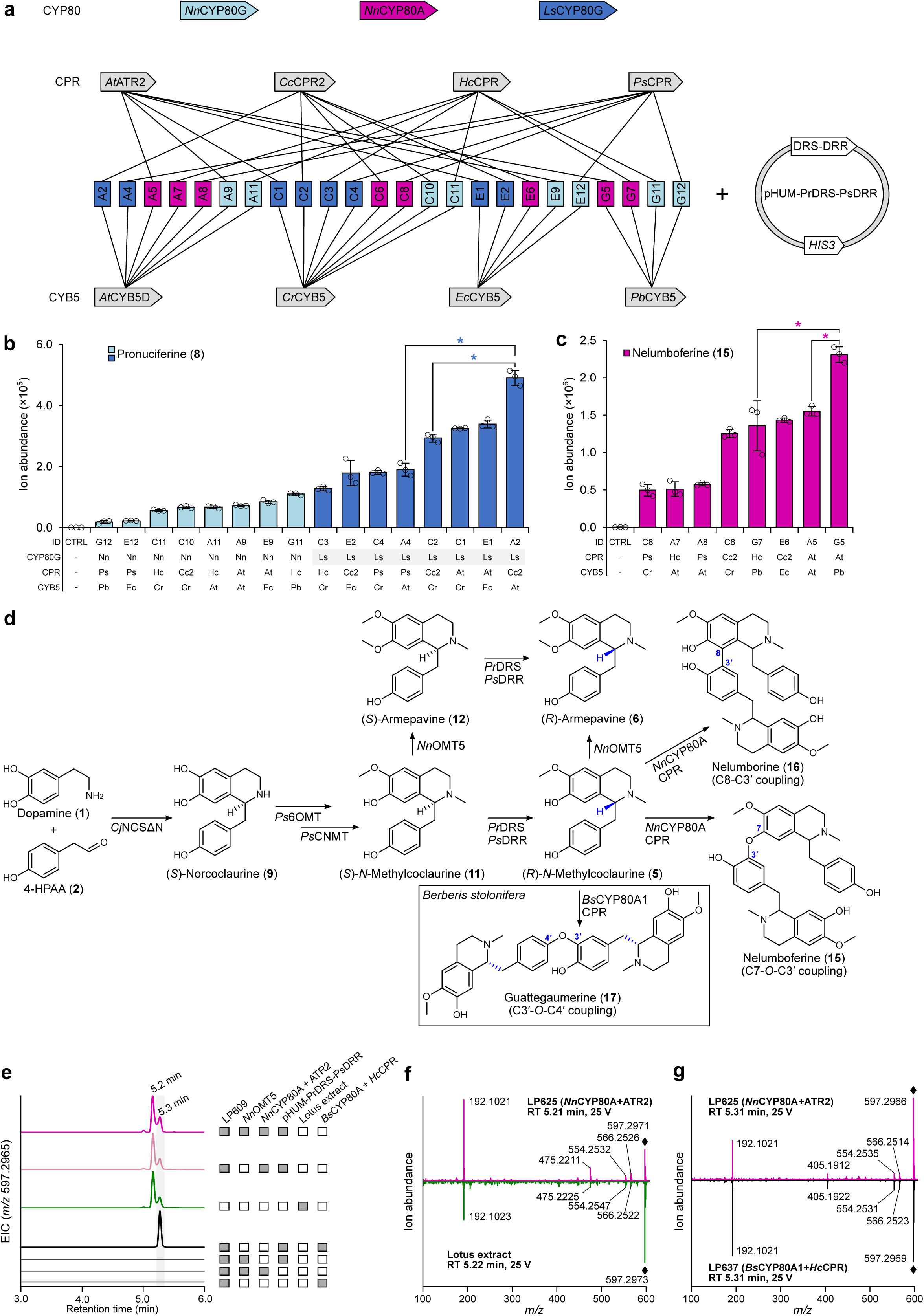
Combinatorial CYP80-CPR-CYB5 library for improved pronuciferine synthesis and characterization of *Nn*CYP80A. **a,** Design of a partial 24-member CYP80-CPR-CYB5 combinatorial strain library. Two CYP80G variants (*Nn*CYP80G and *Ls*CYP80G) and CYP80A from *N. nucifera* (*Nn*CYP80A) were paired with different combinations of plant cytochrome P450 reductases (CPRs) and cytochrome *b5* (CYB5s) orthologs. The 24-strain library was assayed with and without pHUM-PrDRS-PsDRR, which expresses *PrDRS* and *PsDRR* genes for stereochemical inversion of BIA metabolites. Abbreviations: At, *Arabidopsis thaliana*; Cc2, *Corydalis cheilanthifolia* CPR2; Cr, *Catharanthus roseus*; Ec, *Eschscholzia californica*; Hc, *Hypericum calycinum*; *Pb*, *Papaver bracteatum*; *Ps*, *P. somniferum*. **b,** Pronuciferine production by strains harboring *Nn*CYP80G or *Ls*CYP80G, pHUM-PrDRS-PsDRR, and different combinations of CPR and CYB5. Asterisk (*) denotes a significant increase or decrease (*P* < 0.05) in pronuciferine ion abundance between the designated strains. All strains exhibited a significant increase (*P* < 0.05) in pronuciferine ion abundance relative to the parent strain lacking a CYP80G, CPR, and CYB5. **c,** bis-BIA synthesis by strains harboring *Nn*CYP80A, pHUM- PrDRS-PsDRR, and different combinations of CPR and CYB5. Asterisk (*) denotes a significant increase or decrease (*P* < 0.05) in bis-BIA ion abundance between the designated strains. All strains exhibited a significant increase (*P* < 0.05) in bis-BIA ion abundance relative to the parent strain lacking *Nn*CYP80A, CPR, and CYB5. **d,** Proposed pathways to lotus bis-BIA alkaloids in engineered yeast. *Nn*CYP80A catalyzes the stereospecific formation of nelumborine (C8-C3ʹ) or nelumboferine (C7-*O*-C3ʹ) from *N*-methylcoclaurine via distinct phenol coupling routes. Guattegaumerine biosynthesis in *Berberis stolonifera* is shown (boxed), which involves C3ʹ-*O*- C4ʹ coupling of two units of (*R*)-*N*-methylcoclaurine by *Bs*CYP80A1. **e,** Ion-extracted LC–MS chromatograms of engineered yeast strains harboring *Nn*CYP80A or *Bs*CYP80A1, pHUM- PrDRS-PsDRR, and a CPR. The bis-BIA product from *Nn*CYP80A-producing strains was compared against commercial lotus extract. **f,** MS/MS fragmentation of a bis-BIA product (RT ∼5.2 min) from sacred lotus extract and strain LP625 containing *Nn*CYP80A and *At*ATR2. **g,** MS/MS fragmentation of bis-BIA products (RT ∼5.3 min) from strain LP625 (*Nn*CYP80A+*At*ATR2) and LP637 (*Bs*CYP80A1+*Hc*CPR). The 597 → 192 transition is the major product observed upon fragmentation of authentic guattegaumerine standard (Payne et al., 2021).

Expression of *PrDRS* and *PsDRR* in strains containing *Nn*CYP80A and different combinations of CPR and CYB5 triggered the formation of a new LC-MS metabolite peak ([M + H]^+^ = 597.2965 Da) consistent with a high molecular weight bis-BIA product (**Fig. 3c,d**). Selection of both CPR and CYB5 variant had a significant effect on production of the tentative bis-BIA product. The bis-BIA metabolite(s) eluted as a double peak and its formation required *Nn*CYP80A as well as *Pr*DRS and *Ps*DRR (**Fig. 3e**; dark pink trace), indicating a requirement for (*R*)-substrates.

At least seven bis-BIAs have been isolated from *N. nucifera* (Menéndez-Perdomo and Facchini, 2018), of which four (nelumborine, nelumboferine, 6-hydroxynorisoliensinine, and *N*- norisoliensinine) have a molecular weight and chemical formula (C_36_H_40_N_2_O_6_) consistent with the metabolite peak generated by yeast strains harboring *Nn*CYP80A, *Pr*DRS and *Ps*DRR. Of these tentative products, nelumboferine (**15**) and nelumborine (**16**) possesses a methylation profile consistent with methyltransferases produced by our combinatorial library, as 6- hydroxynorisoliensinine and *N*-norisoliensinine lack methyl groups furnished by *Ps*6OMT and *Ps*CNMT, respectively. Nelumboferine and nelumborine both derive from the coupling of two monomers of *N*-methylcoclaurine (Itoh et al., 2011), indicating that *Nn*OMT5 is not involved in their formation (**Fig. 3d**). To clarify the role of *Nn*OMT5 in the synthesis of the tentative bis-BIA metabolite, we deleted *NnOMT5* from the best performing *Nn*CYP80A strain harboring *At*ATR2 and *Pb*CYB5. In agreement with the presumed routes to nelumboferine and nelumborine in *N. nucifera*, deletion of *NnOMT5* had no effect on synthesis of the bis-BIA metabolite (**Fig. 3e**; light pink trace).

To further probe the identity of the tentative bis-BIA metabolite, we analyzed sacred lotus extracts for the presence of high molecular weight bis-BIA metabolites. We identified a metabolite peak with the same exact mass ([M + H]^+^ = 597.2965 Da) and retention time in lotus leaf extract as the tentative bis-BIA product synthesized by yeast strains harboring *Nn*CYP80A, *Pr*DRS and *Ps*DRR (**Fig. 3e**; green trace). In line with yeast strains expressing *NnCYP80A, PrDRS* and *PsDRR*, the lotus bis-BIA metabolite eluted as a double peak in commercial lotus extract. Targeted fragmentation of the tentative bis-BIA product from lotus extract and yeast supernatants yielded several common fragment ions, notably [M + H]^+^ = 192.1, 475.2, 554.2, and 566.2 Da (**Fig. 3f**).

We also screened the CYP80-CPR-CYB5 combinatorial strain library in the absence of a DRS-DRR pair to search for tentative products arising from exclusively (*S*)-metabolites. In agreement with known stereochemistry of lotus alkaloids (Menéndez-Perdomo and Facchini, 2018), no tentative bis-BIA or proaporphine metabolite peaks were identified, confirming an inability of *Nn*CYP80A, *Nn*CYP80G, and *Ls*CYP80G to turnover substrates possessing exclusively (*S*)-configurations.

Guattegaumerine (**17**) is a bis-BIA metabolite synthesized by *Berberis stolonifera*. Both guattegaumerine and nelumboferine derive from two units of (*R*)-*N*-methylcoclaurine, yet guattegaumerine derives from 3ʹ-*O*-4ʹ phenol coupling, while nelumboferine arises from 7-*O*-3ʹ coupling (**Fig. 3d**). Recently, yeast was engineered to synthesize guattegaumerine from (*R*)-*N*- methylcoclaurine by employing CYP80A1 from *B. stolonifera* (*Bs*CYP80A1), the product of which was validated using authentic guattegaumerine standard (Payne et al., 2021). Because guattegaumerine, nelumboferine, and nelumborine are structural isomers (C_36_H_40_N_2_O_6_) generated through different phenol coupling reactions, we reasoned that our *Nn*CYP80A-derived bis-BIA product would yield a similar LC-MS retention time and MS/MS fragmentation spectra to that of *Bs*CYP80A1-derived guattegaumerine. To first demonstrate synthesis of guattegaumerine, we paired *Bs*CYP80A1 with four CPR variants in our (*R*)-*N*-methylcoclaurine host lacking *NnOMT5*. All *Bs*CYP80A1 strains facilitated the synthesis of a new LC-MS peak with the same exact mass as guattegaumerine ([M + H]^+^ = 597.2965 Da) (**Fig. 3e**; black trace). Pairing of *Bs*CYP80A1 with *Hc*CPR yielded approximately 45-fold more guattegaumerine than an isogenic strain harboring *Cc*CPR2 (**Supplementary Fig. 2**). Production of guattegaumerine required expression of *PrDRS* and *PsDRR*, corroborating the requirement for (*R*)-*N*-methylcoclaurine by *Bs*CYP80A1 (Payne et al., 2021). In contrast to the *Nn*CYP80A-derived bis-BIA product, guattegaumerine eluted as a single LC-MS peak, yet with the same retention time as the *Nn*CYP80A product, presumably due to the different phenol coupling reactions involved in the synthesis of guattegaumerine, nelumboferine, and nelumborine. We compared MS/MS fragmentation of *Nn*CYP80A- and *Bs*CYP80A1-derived products (**Fig. 3f**), which yielded the same major 597→192 transition generated by fragmentation of authentic guattegaumerine standard (Payne et al., 2021). Collectively, these data are consistent with a tentative assignment of nelumboferine or nelumborine for the identity of the *Nn*CYP80A-derived bis-BIA product.

### Derivatization of nelumboferine by screening a library of BIA *O*-methyltransferases

In sacred lotus, nelumboferine (**15**) is converted to liensinine (**18**) or isoliensinine (**19**) via methylation by 7OMT or 4ʹOMT, respectively (Itoh et al., 2011). One or both of these compounds are subsequently methylated to give neferine (**20**) (**Fig. 4a**). We screened our in-house library of BIA methyltransferase enzymes for activity on nelumboferine. In total we introduced 66 putative *O*-methyltransferases from 19 BIA-producing plant species to our top-performing *Nn*CYP80A strain (**Fig. 4b**). Our search included seven predicted *O*-methyltransferases from *N. nucifera* (*Nn*6OMT1-4 and *Nn*7OMT1-3).

**Figure 4.**
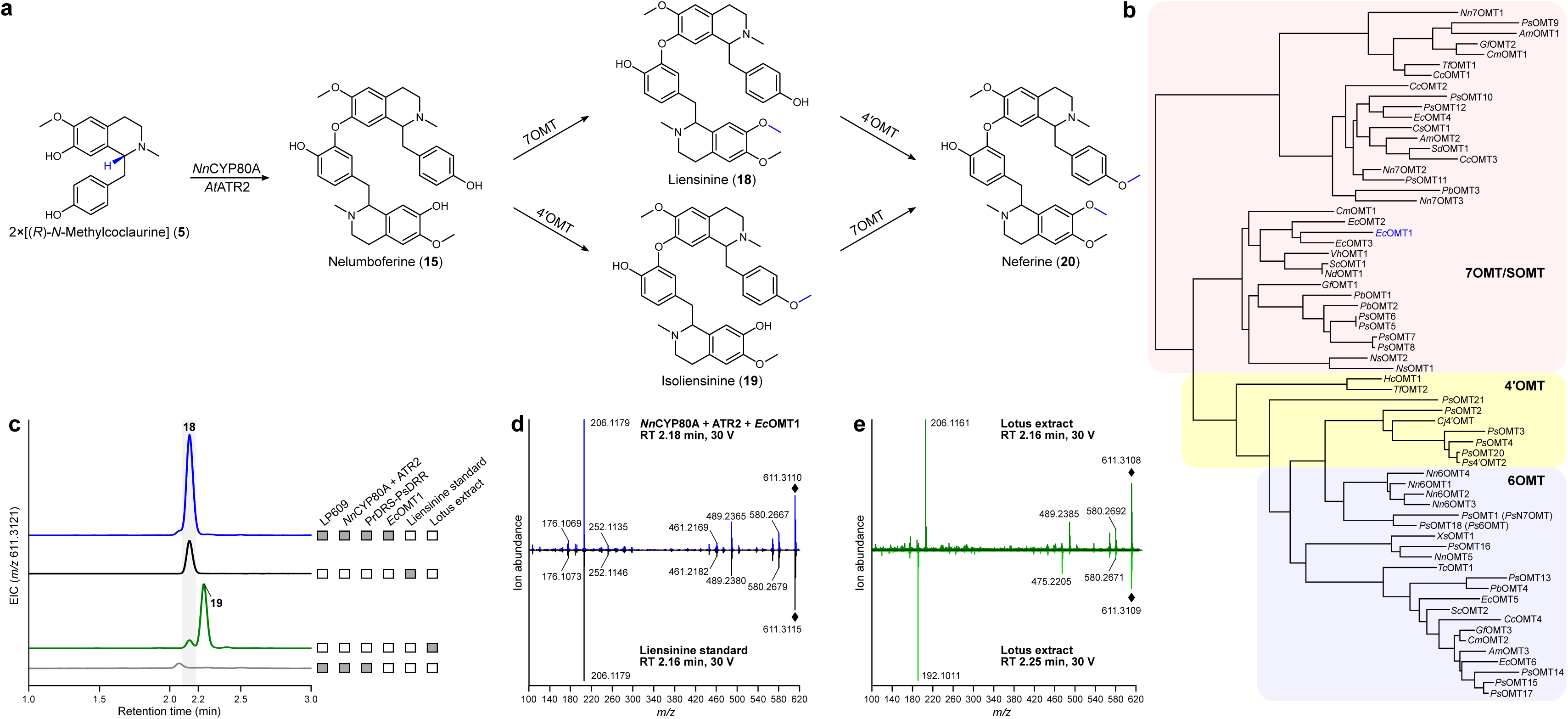
Screening diverse plant *O*-methyltransferases for activity on nelumboferine. **a,** Proposed neferine biosynthesis pathways in *N. nucifera*. Neferine is synthesized via convergent routes from liensinine or isoliensinine, both of which are presumed to arise from nelumboferine. **b,** Protein phylogenetic tree of 66 *O*-methyltransferase orthologs from various plant species. Abbreviations: *Am*, *Argemone mexicana*; *Cc*, *Corydalis cheilanthifolia*; *Cj*, *Coptis japonica*; *Cm*, *Chelidonium majus*; *Cs*, *Corydalis saxicola*; *Ec*, *Eschscholzia californica*; *Gf*, *Glaucium flavum*; *Hc*, *Hypericum calycinum*; *Nd*, *Nandina domestica*; *Nn*, *Nelumbo nucifera*; *Ns*, *Nigella sativa*; *Pb*, *Papaver bracteatum*; *Ps*, *Papaver somniferum*; *Sc*, *Sanguinaria canadensis*; *Sd*, *Stylophorum diphyllum*; *Tc*, *Tinospora cordifolia*; *Tf*, *Thalictrum flavum*; *Vh*, *Vancouveria hexandra*; Xs, *Xanthoriza simplicissima.* **c,** Ion-extracted LC–MS chromatograms of engineered yeast strains harboring *Nn*CYP80A, *At*ATR2, *Pr*DRS-*Ps*DRR, and *Ec*OMT1 from *Eschscholzia californica*. Retention time of yeast-derived liensinine (dark blue) was compared against authentic liensinine standard (black) and sacred lotus extract (green). **d,** MS/MS fragmentation of authentic liensinine standard and a yeast strain engineered for liensinine synthesis (*Nn*CYP80A, ATR2, *Pr*DRS, *Ps*DRR, and *Ec*OMT1). **e,** MS/MS fragmentation of tentative liensinine (RT = 2.16 min; dark green) and isoliensinine (RT = 2.25 min; light green) products from commercial lotus extract. Liensinine (611 → 206) and isoliensinine (611 → 192) yield different major fragment ions based on a prior study (Zhou et al., 2007).

We analyzed cultures of methyltransferase-expressing strains for new peaks consistent with single ([M + H]^+^ = 611.3121 Da) or double methylation ([M + H]^+^ = 625.3278 Da) of the *Nn*CYP80A-derived bis-BIA product ([M + H]^+^ = 597.2965 Da). One *O*-methyltransferase candidate (*Ec*OMT1 from *Eschscholzia californica*) generated a peak ([M + H]^+^ = 611.3121 Da) consistent with single methylation (**Fig. 4c**). Based on a prior study, *Ec*OMT1 is a 7OMT that catalyzes the conversion of reticuline to laudanine (Fujii et al., 2007). In line with its reported activity, *Ec*OMT1 clusters with other enzymes exhibiting confirmed or predicted 7OMT activity (**Fig. 4b**), including *Ec*OMT2, a scoulerine 9OMT (SOMT) that also possesses 7OMT activity on reticuline (Purwanto et al., 2017). We analyzed the active site of *Ec*OMT1 using AlphaFold (Jumper et al., 2021; Varadi et al., 2022) and observed a unique Gly256 residue that is predicted to form a hydrogen bond with the His259 catalytic residue (**Supplementary Fig. 3**) conserved across all BIA OMTs (Morris and Facchini, 2019). Gly256 is substituted with a bulky Trp residue in other closely related BIA OMTs within the *Ec*OMT1- and *Ec*OMT2-containing 7OMT clade, which might hinder binding of large bis-BIA substrates. Similarly, Gly308 of *Ec*OMT1 is believed to play a role in the alleviation of substrate steric hindrance (Morris and Facchini, 2019) and related 7OMTs harbor a bulkier Ile, Leu, or Val residue in this position.

Based on the reported 7OMT activity of *Ec*OMT1, the identity of the new methylated LC-MS product is expected to be liensinine (C_37_H_42_N_2_O_6_) (**Fig. 4a**), a major bis-BIA in lotus. We obtained authentic liensinine standard, which generated a mass ([M + H]^+^ = 611.3121 Da) and retention time (RT = 2.16 min) in agreement with the LC-MS product derived from *Ec*OMT1 activity on nelumboferine (**Fig. 4c**). We also compared MS/MS fragmentation of the tentative liensinine metabolite peak with authentic liensinine standard, which generated identical fragmentation spectra (**Fig. 4d**). Specifically, the major fragment ion with mass [M + H]^+^ = 206.1 was previously used to differentiate liensinine from its isomer isoliensinine (C_37_H_42_N_2_O_6_) (Zhou et al., 2007), which arises from 4ʹ-*O*-methylation of nelumboferine in place of 7-*O*- methylation. The fragment ion with mass [M + H]^+^ = 206.1 also differentiates liensinine from its nelumboferine precursor (C_36_H_40_N_2_O_6_), which yields a corresponding fragment ion with mass [M + H]^+^ = 192.1 (**Fig. 3f,g**), indicating *O*-methylation of the free tetrahydroisoquinoline moiety of nelumboferine. Importantly, synthesis of liensinine by yeast strains harboring *Nn*CYP80A, ATR2, and *Ec*OMT1 confirms that nelumboferine is a product of CYP80A from *N. nucifera*.

We also identified liensinine in commercial lotus extract (RT = 2.16 min), although an additional larger peak (RT = 2.25 min) with the same mass was also observed (**Fig. 4c**). As expected, MS/MS fragmentation of the minor lotus peak yielded the key 611 → 206 transition characteristic of liensinine (**Fig. 4e**; dark green). In contrast, fragmentation of the dominant lotus product (RT = 2.25 min) yielded a major 611 → 192 transition (**Fig. 4e**; light green), which is in agreement with isoliensinine (Zhou et al., 2007).

We also introduced our panel of 66 methyltransferases to our guattegaumerine-producing strain harboring *Bs*CYP80A1 and *Hc*CPR, yet were unable to detect methylated derivatives ([M + H]^+^ = 611.3121 Da or 625.3278 Da). Our screen identified several methyltransferases with tentative 7OMT or 4ʹOMT activities on *N*-methylcoclaurine (**Supplementary Table 1**).

## DISCUSSION

In contrast to most BIA-producing plants, in which the core BIA pathway proceeds through exclusively (*S*)-conformers, sacred lotus synthesizes (*R*)-*N*-methylcoclaurine and (*R*)-armepavine in a dedicated (*R*)-stereochemical path from (*R*)-norcoclaurine. In this study, we designed and implemented a synthetic strategy to flip the stereochemistry of our BIA platform strain by exploiting the unique substrate preference of DRS from common poppy (*Pr*DRS). Pairing of *Pr*DRS with several DRR variants enabled stereochemical inversion of (*S*)-*N*-methylcoclaurine and (*S*)-armepavine via 1,2-dehydro-*N*-methylcoclaurine and 1,2-dehydro-armepavine intermediates. Production of (*R*)-conformers enabled the characterization of *Nn*CYP80G and *Nn*CYP80A as respective stereospecific proaporphine and bis-BIA synthases involved in the committed steps of the aporphine and bis-BIA branches of BIA metabolism in sacred lotus.

Our findings firmly establish roles for *Nn*CYP80G and *Nn*CYP80A in propaporphine and bis-BIA biosynthesis, respectively. Specifically, we show that *Nn*CYP80G is a proaporphine synthase that catalyzes the stereospecific conversion of (*R*)-*N*-methylcoclaurine to glaziovine, which is subsequently methylated to pronuciferine by *Nn*OMT5. Formation of pronuciferine was dependent on implementation of *Pr*DRS-*Ps*DRR or exogenous supplementation of (*R*,*S*)- norcoclaurine, clearly establishing the stereospecificity of *Nn*CYP80G for (*R*)-*N*- methylcoclaurine. Our findings, coupled with the recent discovery and characterization of *Nn*OMT5 (Menéndez-Perdomo and Facchini, 2020), the dedicated 7OMT responsible for armepavine and pronuciferine production, have unlocked the aporphine branch of BIA biosynthesis in sacred lotus. Our data is in agreement with the phenol coupling mechanism of aporphine formation set forth by Barton and Cohen in 1957, in which the authors predicted the existence of proaporphine (dienone) intermediates prior to the isolation of any such metabolites from natural sources (Barton and Cohen, 1957). Soon after, many proaporphine natural products were isolated from plants, including pronuciferine, mecambrine, stepharine, crotonosine, and glaziovine (Stuart and Cava, 1968). According to the phenol coupling mechanism, 7-*O*- methylated substrates, such as isococlaurine and armepavine, are not converted into proaporphine structures, which was validated using *in vivo* tracer studies (Barton et al., 1967; Haynes et al., 1965). Our findings from yeast supplementation assays are in agreement with this general mechanism, as feeding (*R*,*S*)-norcoclaurine, but not (*R*,*S*)-armepavine, facilitated synthesis of pronuciferine, suggesting that *Nn*CYP80G acts on a substrate upstream of armepavine. By deleting *Nn*OMT5, we showed that *Nn*CYP80G catalyzes the formation of glaziovine from (*R*)-*N*-methylcoclaurine based on exogenous supplementation of (*R*,*S*)- norcoclaurine. While glaziovine is not a major constituent of sacred lotus (Menéndez-Perdomo and Facchini, 2018), several recent studies have reported its isolation from seeds of *N. nucifera* (Bishayee et al., 2022; Cao et al., 2022; Yu et al., 2022). In contrast, glaziovine is a major alkaloid in *Ocotea glaziovii* and other select species of Lauraceae and Annonaceae (Agnès et al., 2022). Our demonstration of glaziovine synthesis coupled with the inability of *Nn*CYP80G to act on exogenous (*R*,*S*)-armepavine confirms that proaporphine synthesis precedes 7-*O*-methylation in agreement with the proposed phenol coupling mechanism (Barton and Cohen, 1957). Although a recent study failed to demonstrate *Nn*OMT5 activity on aporphine substrates and thus postulated that 7OMT precedes aporphine synthesis (Menéndez-Perdomo and Facchini, 2020), glaziovine and other proaporphine metabolites were not assayed. We show that *Nn*OMT5 exhibits appreciable activity on both 1-benzylisoquinoline (*N*-methylcoclaurine) and proaporphine substrates (glaziovine), indicating that *N*-methylcoclaurine is the major branchpoint metabolite in sacred lotus and 7-*O*-methylation acts as a blocking group to limit flux to the aporphine pathway.

Nuciferine, a major alkaloid in *N. nucifera* and a metabolite with potential anti-cancer properties, possesses the same methylation pattern as pronuciferine, indicating that a hitherto unidentified enzyme is responsible for conversion of pronuciferine to nuciferine. To identify the unknown activity, we implemented 10 lotus metabolic enzymes with expression profiles similar to *Nn*CYP80G and other core BIA pathway enzymes. However, we were unable to identify the elusive missing enzyme responsible for the conversion of pronuciferine to nuciferine. More sophisticated enzyme discovery strategies, such as those employed in the recent elucidation of colchicine and strychnine biosynthetic pathways (Hong et al., 2022; Nett et al., 2020), will be required to complete the lotus pathway to nuciferine.

*Nn*CYP80A was previously implicated in bis-BIA biosynthesis, while other studies have proposed it to be an alternative isoform of *Nn*CYP80G (Menéndez-Perdomo and Facchini, 2018). We leveraged our artificial stereochemical inversion strategy to clarify the role of *Nn*CYP80A in sacred lotus metabolism. We show that *Nn*CYP80A is the dedicated bis-BIA synthase involved in the committed step of the *N. nucifera* bis-BIA biosynthesis pathway. To date seven high molecular weight bis-BIA metabolites have been isolated from sacred lotus, in addition to an unusual tri-BIA (neoliensinine) (Menéndez-Perdomo and Facchini, 2018). Of the characterized lotus bis-BIAs, only nelumboferine and nelumborine possess the exact mass ([M + H]^+^ = 597.2965 Da; C_36_H_40_N_2_O_6_) and methylation profile consistent with the *Nn*CYP80A- catalyzed metabolite. Nelumborine and nelumboferine possess the same methylation pattern yet derive from distinct phenol coupling reactions. Whereas nelumboferine and other lotus bis-BIAs derive from a C7-*O*-C3ʹ phenol coupling, nelumborine arises from an unusual C8-C3ʹ coupling. We showed that formation of the *Nn*CYP80A bis-BIA product was independent of *Nn*OMT5, yet required the presence of *Pr*DRS-*Ps*DRR, implicating involvement of (*R*)-*N*-methylcoclaurine in its synthesis. Consistent with this observation, nelumboferine arises from two units of (*R*)-*N*- methylcoclaurine (Itoh et al., 2011; Nishimura et al., 2013), while to our knowledge the stereochemistry of *N*-methylcoclaurine monomers in the synthesis of nelumborine has not been deduced.

Based on the methylation profile of BIAs isolated from *N. nucifera*, nelumboferine gives rise to neferine, the fully methylated bis-BIA end product in lotus, via convergent routes from liensinine or isoliensinine (Itoh et al., 2011) involving unidentified 7OMT and 4ʹOMT enzymes. We introduced our library of 66 plant methyltransferases into our top bis-BIA-producing strain, including seven predicted OMTs from *N. nucifera*, and identified *Ec*OMT1 from California poppy (*Eschscholzia californica*) as a promiscuous 7OMT with activity on nelumboferine. To our knowledge, this is the first plant OMT capable of methylating bis-BIAs and we identified two unique *Ec*OMT1 active site residues (Gly256 and Gly308) that are not conserved across closely related plant 7OMT orthologs, such as *Ec*OMT2. These residues might play a role in positioning the nelumboferine substrate or alleviating steric hindrance of bulky bis-BIAs (Morris and Facchini, 2019). Interestingly, *Ec*OMT1 did not exhibit 7OMT activity on guattegaumerine, which likely reflects the distinct conformations adopted by guattegaumerine and nelumboferine isomers. Implementation of *Ec*OMT1 in a strain containing *Nn*CYP80A, ATR2, *Pr*DRS, and *Ps*DRR facilitated synthesis of liensinine, which was confirmed using an authentic liensinine standard. Liensinine is a major bis-BIA metabolite in sacred lotus with promising anti-cancer activity (Zhou et al., 2015) and our findings establish an artificial *de novo* path for microbial overproduction of liensinine. While we did not identify the dedicated *N. nucifera* bis-BIA 7OMT involved in liensinine formation, our findings might aid in the identification of the elusive enzyme by using *Ec*OMT1 to query the lotus genome, which encodes at least 60 candidate *O*- methyltransferase genes (Li et al., 2021). Moreover, repeating our screen of diverse plant *O*- methyltransferases might unveil a novel 4ʹOMT variant with activity on liensinine, thus enabling synthesis of neferine, another prospective anti-cancer pharmaceutical.

Formation of liensinine by the concerted activity of *Nn*CYP80A and *Ec*OMT1 further confirms nelumboferine as a product of CYP80A activity in *N. nucifera*. We compared *Nn*CYP80A activity to that of *Bs*CYP80A1, a recently characterized bis-BIA synthase from *Berberis stolonifera*. *Bs*CYP80A1 catalyzes the 3ʹ-4ʹ C-O phenol coupling of two units of (*R*)-*N*- methylcoclaurine to give guattegaumerine or one unit each of (*R*)- and (*S*)-*N*-methylcoclaurine, yielding berbamunine (Payne et al., 2021). Products of *Bs*CYP80A1 and *Nn*CYP80A possess the same mass ([M + H]^+^ = 597.2965 Da; C_36_H_40_N_2_O_6_) and expected methylation profile, yet the *Nn*CYP80A product eluted as two distinct LC-MS peaks, while strains harboring *Bs*CYP80A1 produced a single peak. Although we confirmed nelumboferine as a product of *Nn*CYP80A based on conversion to liensinine, it is possible that *Nn*CYP80A is involved in the biosynthesis of both nelumboferine and nelumborine by catalyzing dual coupling reactions (C7-*O*-C3ʹ and C8-C3ʹ, respectively), which might provide justification for the two LC-MS peaks derived from *Nn*CYP80A activity. However, the biosynthetic origin of nelumborine in *N. nucifera* remains unclear.

*NnCYP80A* and *NnCYP80G* genes are adjacent in the genome of *N. nucifera*, suggesting a tandem duplication event. Despite different roles in BIA metabolism, the respective *Nn*CYP80A and *Nn*CYP80G enzymes share a significantly higher degree of sequence identity (80% amino acid identity) than *Bs*CYP80A1 and *Nn*CYP80A bis-BIA synthases (45% amino acid identity), both of which catalyze the coupling of two units of (*R*)-*N*-methylcoclaurine. Surprisingly, based on amino acid sequence, *Bs*CYP80A1 is more closely related to lotus *Nn*CYP80G aporphine synthase than the analogous lotus *Nn*CYP80A bis-BIA synthase. Owing to their similarity, earlier studies assumed that *Nn*CYP80G and *Nn*CYP80A (also referred to as *Nn*CYP80Q1 and *Nn*CYP80Q2) (Nelson and Schuler, 2013) both encode bis-BIA synthases. Instead, our findings suggest that following gene duplication, *Nn*CYP80A and *Nn*CYP80G diverged in activity. These observations suggest that it might be possible to switch *Nn*CYP80G to a bis-BIA synthase or *Nn*CYP80A to an aporphine synthase with a potentially small number of amino acid changes. Our characterization of *Nn*CYP80A and *Nn*CYP80G adds to the diverse collection of plant CYP80 variants, including corytuberine synthase (CYP80G2), (*S*)-*N*- methylcoclaurine 3ʹ-hydroxylase (CYP80B1), and CYP80F1 involved in littorine rearrangement, with key roles in benzylisoquinoline and tropane alkaloid metabolism. An in-depth structural analysis of such broad CYP80 enzymes is likely to shed light on key residues involved in substrate binding and catalytic mechanism.

In addition to *Nn*CYP80A and *Bs*CYP80A1 variants, Asian moonseed (*Menispermum dauricum*) and Canadian moonseed (*M. canadense*) synthesize dauricine, a potential anti-cancer bis-BIA that arises from the C3ʹ-*O*-C4ʹ coupling of two units of (*R*)-armepavine. A previous study screened five plant CYP80A orthologs in an (*R*)-armepavine-producing strain, yet did not observe production of dauricine (Payne et al., 2021). We attempted to devise an artificial route to dauricine through direct methylation of guattegaumerine by screening our library of 66 plant methyltransferases in a strain harboring *Bs*CYP80A1 and *Hc*CPR. However, we were also unsuccessful in synthesizing dauricine or methylated derivatives of guattegaumerine. A more exhaustive screen of diverse plant CYP80A orthologs on stereoisomers of both *N*- methylcoclaurine and armepavine might unveil new coupling activities, such as the C3ʹ-*O*-C4ʹ coupling route of (*R*)-armepavine in the biosynthesis of dauricine.

To the best of our knowledge, a dedicated (*R*)- or (*R*,*S*)-yielding NCS has yet to be identified from sacred lotus, which prompted our artificial route involving *Pr*DRS and *Ps*DRR (REPI). While our stereochemical inversion approach enabled the use of the highly active (*S*)- specific *Cj*NCSΔN variant (Grewal et al., 2021; Pyne et al., 2020), the native lotus route from (*R*)-norcoclaurine bypasses the requirement of DRS and DRR enzymes. Thus, it is unclear if our stereochemical inversion path is more efficient than the native route when reconstructed in engineered yeast. Screening all five of the predicted lotus NCS variants (Menéndez-Perdomo and Facchini, 2018) might lead to the discovery of the presumed (*R*)-specific isoform. Relatedly, we showed that two lotus CNMT variants (*Nn*CNMT1 and *Nn*CNMT3) lacked activity in a *Cj*NCSΔN strain, suggesting that these enzymes might be specific for (*R*)-substrates or remaining unidentified lotus methyltransferases are involved in (*R*)-*N*-methylcoclaurine synthesis. Consequently, several core enzymes involved in the (*R*)-specific BIA pathway remain unidentified in *N. nucifera*.

Finally, our study highlights the utility of high-titer microbial strains in plant enzyme discovery and pathway elucidation. BIA output from our (*S*)-norcoclaurine strain exceeds 1.5 g/L (Pyne et al., 2020), which provides an ideal background for enzyme characterization, particularly those with poor catalytic properties or weak off-target activities. By repurposing our BIA platform, we achieved *de novo* synthesis of 152 mg/L armepavine, a 14-fold improvement over a previous study (11.1 mg/L armepavine) in which L-DOPA precursor supplementation was employed to boost BIA synthesis (Payne et al., 2021). Relatedly, guattegaumerine output varied by a factor of 45 across all four of our *Bs*CYP80A1-CPR strains, while an earlier study was unable to detect guattegaumerine in two out of four CPR pairings (Payne et al., 2021). We also detected pronuciferine or bis-BIA synthesis from all 24 strains of our combinatorial CYP80- CPR-CYB5 library, again showcasing the utility of our high titer BIA platform in the discovery and elucidation of plant secondary metabolic pathways. Our characterization of both *Nn*CYP80A and *Nn*CYP80G herein will enable the exploration and overproduction of potential aporphine and bis-BIA therapeutics from sacred lotus, such as nuciferine, liensinine, and neferine.

## Acknowledgements

We thank the Concordia Genome Foundry for assistance with combinatorial strain construction and Marcos DiFalco and Lauren Narcross for assistance with LC-MS analyses. This study was financially supported by an NSERC-Industrial Biocatalysis Network (IBN) grant and an NSERC Discovery grant. M.E.P. was supported by an NSERC Postdoctoral Fellowship. V.J.J.M. is supported by a Concordia University Research Chair.

## Author contributions

M.E.P. designed the research and performed the experiments with supervision from V.J.J.M.

N.D.G. assisted with combinatorial pathway assembly and LC-MS analyses. M.E.P. and V.J.J.M. wrote the manuscript.

## Competing financial interests

The authors declare no competing financial interests.

## Additional information

Supplementary information is available in the online version of the paper. Correspondence and requests for materials should be addressed to M.E.P. and V.J.J.M.

## Data availability

Data supporting the findings of this work are available within the paper and its SI Appendix. Source data for the figures in this paper and the SI Appendix are provided in Dataset S1.

## METHODS

### Chemicals and reagents

General chemicals and reagents were purchased commercially, including dopamine (Sigma-Aldrich), (*S*)-norcoclaurine (Toronto Research Chemicals), 4- hydroxyphenylacetaldehyde (Toronto Research Chemicals), liensinine (MedChemExpress), and (*R*,*S*)-armepavine (MuseChem), unless stated otherwise. Lotus leaf extract was purchased from Amazon (Plum Flower Brand He Ye) or eBay (Hawaii Pharm Pink Lotus herbal extract).

### Plasmids, strains, and growth media

The quadruple auxotrophic *S. cerevisiae* strain BY4741 (*MATa his3Δ1 leu2Δ0 met15Δ0 ura3Δ0*) is the origin of all engineered strains in this study. Strain LP478 was utilized as the basis for this work (Pyne et al., 2020). Cultures were grown in YPD (10 g/L Bacto Yeast Extract, 20 g/L Bacto peptone, 20 g/L glucose) or synthetic complete (SC) medium [6.7 g/L Difco Yeast Nitrogen Base (YNB) without amino acids, 1.62-1.92 g/L Drop-out Medium Supplements (Millipore-Sigma) minus appropriate amino acids, 20 g/L glucose). Recombinant cells were selected on YPD agar containing 400 μg/mL G418, 200 μg/mL hygromycin B, or 200 μg/mL of each antibiotic. Selection using auxotrophic markers was performed in SC medium lacking histidine. Oligonucleotides utilized in this study are listed in Dataset S2. Plasmids and strains employed in this work are provided in **Supplementary Tables 2 and 3**, respectively.

### Yeast strain construction

Genetic modifications to yeast were made via CRISPR-Cas9-mediated genomic integration (Horwitz et al., 2015; Ryan et al., 2014) and *in vivo* DNA assembly (Shao et al., 2009). Cas9 and gRNA were delivered to yeast using pCas-G418 (ref. (Ryan et al., 2014)) or pCas-Hyg (ref. (Bean et al., 2022)). Linear gRNA cassettes were retargeted by PCR and assembled using *in vivo* gap repair with a BsaI- and NotI-digested pCas backbone (Bean et al., 2022). Roughly 150 ng of digested pCas was combined with 300 ng of linear gRNA cassette and 500-1,000 ng of total repair DNA in a standard 50 μL lithium acetate transformation. Cells were heat-shocked at 42 °C for 30 minutes, recovered for 16 hours, and plated onto yeast agar plates containing appropriate antibiotics. Both linear pCas-G418 and pCas-Hyg plasmids were used for complex multi-part DNA assemblies. The CYP80-CPR-CYB5 combinatorial library was assembled using the Echo 550 acoustic liquid handler. Design of gRNAs was performed by selecting top hits common to both CCTop (Labuhn et al., 2017) and a yeast CRISPRi web tool (Smith et al., 2016). Chromosomal loci for DNA integration were selected from previous studies (Bai Flagfeldt et al., 2009; Bourgeois et al., 2018; Mikkelsen et al., 2012; Reider Apel et al., 2016). Gene expression cassettes and genomic integration sites utilized in this work are listed in **Supplementary Tables 4 and 5**, respectively. Synthetic DNAs employed in this study are provided in **Dataset S3**. The yeast *GRE2* gene was replaced with a synthetic DNA landing pad (LP5.T3) possessing a unique Cas9 target site (T3).

### Microtiter plate assay for BIA quantification

Colonies were picked in triplicate into 0.5 mL of 2× SCS medium (13.4 g/L Difco Yeast Nitrogen Base (YNB) without amino acids, 2× Drop-out Medium Supplements (Millipore-Sigma), 40 g/L sucrose) within 96-well deep well plates. Following 24 h of growth, cultures were back-diluted 50× into 0.5 mL of fresh 2× SCS medium in 96-well deep well plates. Cultures were grown at 30 °C with shaking at 350 rpm. Following 48-72 h of growth, OD_600_ measurements were taken, and culture broth was stored at-20 °C for subsequent analysis by LC-MS.

### LC-MS and chiral analysis of BIA metabolites

BIA metabolites were extracted from culture broth containing cells and growth medium. Seven μL of culture broth was combined with 27 μL of 100% acetonitrile (ACN). Following five minutes of shaking at room temperature, 146 μL of 0.12% formic acid was added to give a final concentration of 15% ACN and 0.1% formic acid.

Samples were diluted an additional 10-fold in 15% ACN and 0.1% formic acid. Samples were centrifuged at 4,000 RCF and 10 μL of extracted culture supernatant was separated on a 1290 Infinity II LC system (Agilent Technologies). Metabolites were separated using the following gradient: 2% B to 10% B from 0-4 min (0.3 mL/min), 10% B to 85% B from 4-6 min (0.3 mL/min), held at 85% B from 6-7 min (0.3 mL/min), 85% B to 2% B from 7-7.1 min (0.3 mL/min), and held at 2% B from 7.1-9 min (0.45 mL/min). Solvent A was 0.1% formic acid in water and solvent B was 0.1% formic acid in 100% ACN. Following separation, eluent was injected into an Agilent 6545 quadrupole time-of-flight MS (QTOF-MS; Agilent Technologies) equipped with a Zorbax Eclipse Plus C18 column (50 × 2.1 mm, 1.8 μm; Agilent Technologies). The sample tray and column compartment were set to 4 °C and 30 °C, respectively. The sheath gas flow rate and temperature were adjusted to 10 L/min and 350 °C, respectively, while drying and nebulizing gases were set to 12 L/min and 55 psig, respectively. The drying gas temperature was set to 325 °C. QTOF data was processed and manipulated using Agilent MassHunter Qualitative Analysis software. Norcoclaurine ([M + H]^+^ = 272.1287 Da), coclaurine ([M + H]^+^ = 286.1443 Da), *N*-methylcoclaurine ([M + H]^+^ = 300.1600 Da), armepavine ([M + H]^+^ = 314.1756 Da), 1,2-dehydro-*N*-methylcoclaurine ([M + H]^+^ = 298.1443 Da), and 1,2- dehyroarmepavine ([M + H]^+^ = 312.1600 Da) were identified by exact mass. Norcoclaurine and armepavine were quantified using authentic standards. Pronuciferine ([M + H]^+^ = 312.1600 Da), glaziovine ([M + H]^+^ = 298.1443 Da), and nelumboferine ([M + H]^+^ = 597.2965 Da) were identified by retention time, exact mass, and MS/MS fragmentation by comparison to extracts from leaves of *N. nucifera*. Guattegaumerine ([M + H]^+^ = 597.2965 Da) was identified by exact mass and MS/MS fragmentation. Liensinine ([M + H]^+^ = 611.3121 Da) was identified by retention time, exact mass, and MS/MS fragmentation by comparison to an authentic standard. MS/MS fragmentation of glaziovine, pronuciferine, and guattegaumerine were compared against published fragmentation spectra.

Chiral analysis of norcoclaurine and armepavine was performed by modifying an existing method. Culture supernatant from yeast strains grown in deep-well plates were centrifuged to remove debris and diluted 1:50 or 1:100 in H_2_O. Ten μL was loaded onto a SHODEX ORPak CDBS-453 column and separated chromatographically using a 1290 Infinity II HPLC (Agilent Technologies) and the following gradient: 0–18 min, 5% B; 18–19 min, 95% B; 19–24 min, 95% B; 24-25 min, 5% B; 25-40 min, 5% B where Buffer A was 0.1% formic acid and Buffer B was 0.1% formic acid in ACN. Flow rate was 0.18 mL min^−1^ and temperature was held constant at 25 °C. Compounds eluting from the column were analyzed with a 6560 Ion Mobility QTOF mass spectrometer (Agilent Technologies) with the following settings: gas temp, 325 °C; drying gas, 12 L min^−1^; nebulizer 55 psig; sheath gas temp 350 °C; sheath gas flow 10 L min^−1^; capillary voltage 4000 V; nozzle voltage 500 V; fragmentor 400 V. Retention times of (*S*)- and (*R*)- norcoclaurine were established by comparison to an (*S*)-norcoclaurine authentic standard (Toronto Research Chemicals Inc.) and *rac*-norcoclaurine, which was spontaneously condensed from dopamine (Sigma) and 4-HPAA (Toronto Research Chemicals) in a KH_2_PO_4_/ACN buffer (Pesnot et al., 2012). Retention times of (*S*)- and (*R*)-armepavine were established by comparison to authentic (*R,S*)-armepavine standard and (*S*)-armepavine synthesized by a BY4741 yeast strain harboring *Ps*6OMT, *Ps*CNMT, and *Nn*OMT5 (strain LP626) fed with (*S*)-norcoclaurine.

### Statistical analyses

All numerical values are depicted as means ± s.d. Statistical differences between control and engineered strains were assessed via two-tailed Student’s *t*-tests assuming equal variances using Excel (Microsoft) or Prism (GraphPad Software Inc.). In all cases, *P*- values < 0.05 were considered significant.

## Notes

### Competing Interest Statement

The authors have declared no competing interest.

### Summary of Updates

We performed additional analyses and experiments to clarify the substrate preference of *Nn*CYP80G from sacred lotus and show that the enzyme converts (*R*)-*N*-methylcoclaurine to glaziovine. We also performed a feeding experiment to disprove our initial conclusion that *Nn*CYP80G converts (*R*)-armepavine to pronuciferine.

