## Supplemental Information for "Pathway elucidation and microbial synthesis of proaporphine and bis-benzylisoquinoline alkaloids from sacred lotus (*Nelumbo nucifera*)"

### Supplementary Results

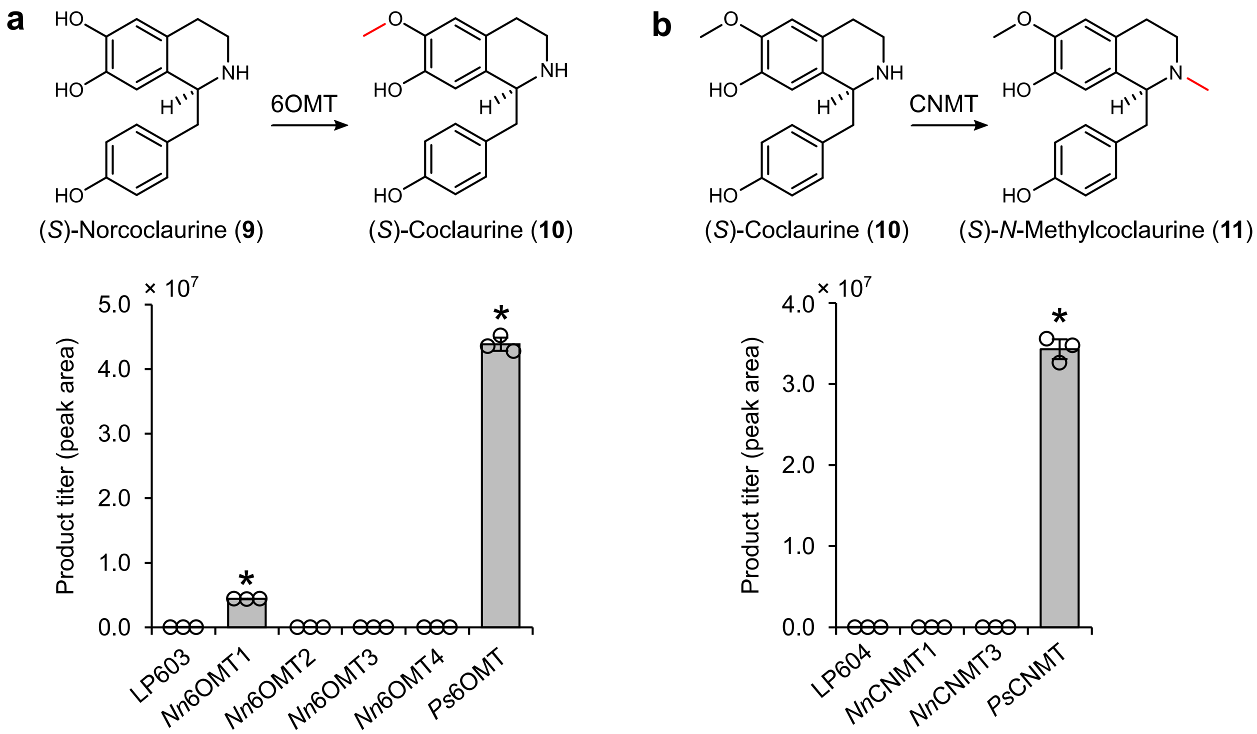


**Supplementary Figure 1. Comparison of core BIA pathway methyltransferases from opium poppy (*Papaver somniferum*) and sacred lotus (*Nelumbo nucifera*). (a)** Screening 6OMT variants for methylation of (*S*)-norcoclaurine to (*S*)-coclaurine. 6OMT isoforms from *N. nucifera* (*Nn*6OMT1-*Nn*6OMT4) were compared against *Ps*6OMT from *P. somniferum* for production of (*S*)-coclaurine in strain LP603. (**b**) Screening CNMT variants for methylation of (*S*)-coclaurine to (*S*)-*N*-methylcoclaurine. CNMT isoforms from *N. nucifera* (*Nn*CNMT1 and *Nn*CNMT3) were compared against *Ps*CNMT from *P. somniferum* for production of (*S*)-*N*-methylcoclaurine in strain LP604 expressing *Ps6OMT*. Asterisk (*) denotes a significant increase ($P < 0.05$) in titer relative to the parent strain. Statistical differences between control and derivative strains were tested using two-tailed Student's *t*-test. Error bars represent s.d. of three biological replicates.

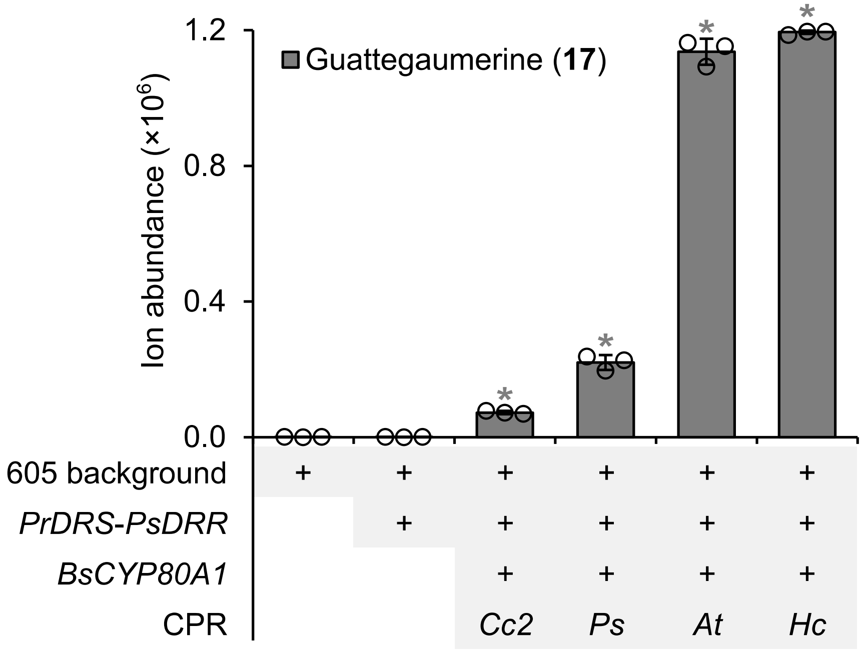


**Supplementary Figure 2. Comparison of guattegaumerine production by strains harboring *Bs*CYP80A1 and different plant CPR variants.** Asterisk (*) denotes a significant increase or decrease ($P < 0.05$) in ion abundance relative to the precursor strain. Error bars represent the mean ± s.d. of $n = 3$ independent biological samples. Statistical differences between control and derivative strains were tested using two-tailed Student's *t*-test. Abbreviations: *At*, *Arabidopsis thaliana*; Cc2, *Corydalis cheilanthifolia* CPR2; Hc, *Hypericum calycinum*; *Ps*, *P. somniferum*.

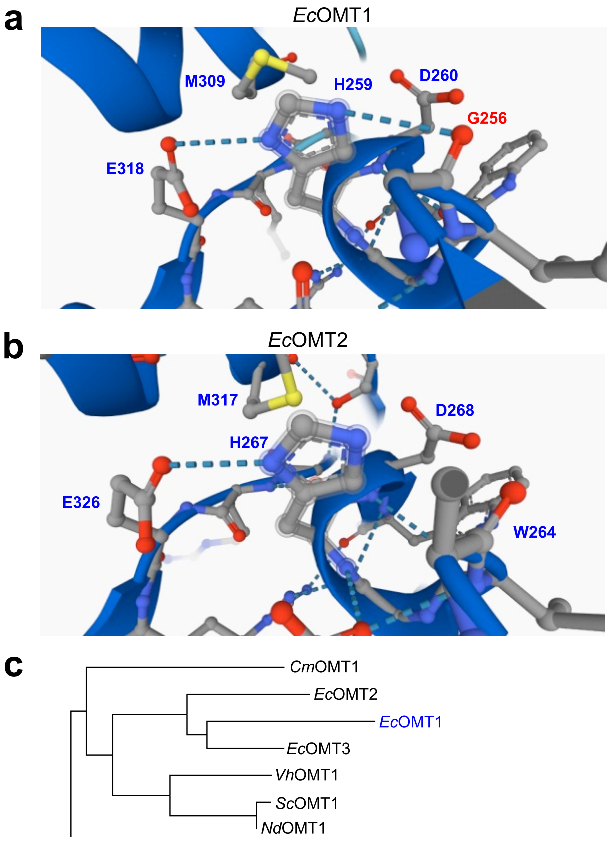


d

```
CmOMT1      VVVESFYNLKEVVLQQASVPFDMTHGVSIFEYAGKDPRMNKVFNEAMGDFSVVAFDEVLK 234
VhOMT1      MLLAPW-HYLSKCVEEGGIAFKKAHGMEIWDFFSSNPELNKVFNEGMACTAKFSVRAILD 179
ScOMT1      MLLAPW-HYLSKVVEEGGVAFAKAHGKEIWELASENEELNKIFNQGMACTSKVTVKAIFS 158
NdOMT1      MLLAPW-HYLSKVVEEGGVAFAKAHGKEIWELASENEELNKVFNQGMACTSKVTVKAIFS 158
EcOMT1      WITDPW-NYLGKCVQEGGSGFVKAHGSDVFKFGSDHPEFFKLFYDGMECSTKVLVHVVLD 182
EcOMT2      WLLDPW-HYLGKCVQEGGFAFAKAHGSEIWKFGSENPEFNKLFSVGMACSSTLVVDAILD 190
EcOMT3      WLLDPW-QYLGKCVQEGGLPFAKAHGSEIWEFGSENPEFNKLFNTGMTCTAKIVVNAILV 167
             :   : :     :::..  *  :** .::.  ... .: *:*  .*   : . .  ::

CmOMT1      VY-NGFLEMKELVDVGGGTGTGIGTSLSNIIAKYPLI-RGINFDLPHVISVAPNYPGVEH 292
VhOMT1      AYKDGFEGIGTLVDLGGGTGAMI----RDIVEEHPHI-KGINFDLPHVVATAPVYPGVEH 234
ScOMT1      VYRDRFEGVRTLVDVGGGTGRMI----GEIVKQHPNVTKGINFDLPHVVATAPEYPGVEH 214
NdOMT1      VYRDRFEGVRTLVDVGGGTGRMI----GEIVKQHPNVTKGINFDLPHVVATAPEYPGVEH 214
EcOMT1      KYQQVFKDVKSIVDVGGGTGMMI----SEIVKNHPNI-KGINFDLPHVVAEAPDYPGVEH 237
EcOMT2      NYHEGFGDLESIVDVGGAIGTLI----NEIVKKYPHI-RGTNFDLPHVVAEALENPGVAH 245
EcOMT3      KYEEGFNDLRSIVDVGGGTGMMI----GEIVKKYPHI-RGINFDLPHVVAAAPEYPGVSH 222
             * : *  :  :**:**. *  *     :*: ::* : :* *******:: *   *** *

CmOMT1      VAGDMFKELPKAQNILLKWVLHDWDNEPCLKLLKSCWNALPE-GGKVIVIEFVVPTKIGD 351
VhOMT1      VGGDMFVAVPNADAVIMKWIMHDWSDGDCVRILKKCREAIPKKTGKVIIVDLMLQPD-GK 293
ScOMT1      VGGDMFAAIPQADAIIMKWIMHDWSDEDCIKILKNCHEAIPKKTGKVIIVDVVLHPE-GE 273
NdOMT1      VGGDMFAAIPQADAIIMKWIMHDWSDEDCIKILKNCHEAIPKKTGKVIIVDVVLHPE-GE 273
EcOMT1      VGGDMFVEIPQADAITMKGILHDWNDDACVRILENCKKAIPK-NGKVIIIDCVLNPD-GD 295
EcOMT2      VGGDMFVEIPSADAVILKWVLHDWNDEDCVKILKNCNKAISN-KGKLIIIECVLKPD-GE 303
EcOMT3      VQGNMFDEIPRADAVIMKWILHDWNDEDSVRILQNCKKAIPK-NGKVIIIDCVLKPD-GK 280
            * *:**  :* *: : :* ::***.:  .:::*:.* :*: :  **:*::: ::  . *.

CmOMT1      NAESFNALTPDLIMMALNPGGKERTLIEYYDLANATGFAKAKAYPISQGFYVIEFHK-- 408
VhOMT1      GIFDNTRLAMDLLMIAHASGGKERTEVEWKKLLKDGGFPRYNIIQLPALQSIIEAYPE- 351
ScOMT1      SLFDGMRLASDLVMIAHVSGGKERTENEWKNLFENGGFPRYKIYQLPTVTSIIEAYPE- 331
NdOMT1      SLFDGTRLAFDLLMIAHASGGKERTENEWKNLLENGGFPRYKIFQLPTVPSIIEAYPE- 331
EcOMT1      DLFDDIKVVSDLGMRVHCSDGKERTEAEWEKLLKKGGFPRYKITHVVTVQSMIEAYPE- 353
EcOMT2      GLFDGLGLAFDLLMIAHSSGGRERTEAEWKKLLKAGGFSRYKITPIKGIPSIIEAYPDI 362
EcOMT3      DLFDETRVVLDLVMIAHTSGGRERTEVEWEKLLKDAGFPRYKITHIPTIQSIIEAYPE- 338
            .  .   :. ** * .   .*:***  *: .* :  ** : :   :     :** :
```

**Supplementary Figure 3. Sequence and structure comparison of *Ec*OMT1 and *Ec*OMT2 active sites.** (**a**) Predicted AlphaFold active site structure of *Ec*OMT1. Conserved residues are shown in blue and a key Gly256 residue unique to *Ec*OMT1 is shown in red. Gly256 is predicted to interact with the highly conserved His259 catalytic residue. (**b**) Predicted AlphaFold active site structure of *Ec*OMT2. Conserved residues are shown in blue. (**c**) Protein phylogenetic tree of plant BIA 7OMT orthologs closely related to *Ec*OMT1. Abbreviations: *Cm*, *Chelidonium majus*; *Ec*, *Eschscholzia californica*; *Nd*, *Nandina domestica*; *Sc*, *Sanguinaria canadensis*; *Vh*, *Vancouveria hexandra*. (**d**) Active site alignment of plant BIA 7OMT orthologs. Conserved active site residues are shown in blue and Gly256 of *Ec*OMT1 is shown in red. Gly308 of *Ec*OMT1 and corresponding residues potentially involved in alleviation of substrate steric hindrance are shown in fuchsia.

**Supplementary Table 1 – Tentative activity assignments of diverse plant *O*-methyltransferases on 1-benzylisoquinolines**

| OMT | Source | Tentative activity | | |
|---|---|---|---|---|
| | | 6OMT | 7OMT | 4′OMT |
| *Am*OMT1 | *Argemone mexicana* | | + | |
| *Am*OMT2 | | | | |
| *Am*OMT3 | | | | |
| *Cc*OMT1 | *Corydalis cheilanthifolia* | | | |
| *Cc*OMT2 | | | | |
| *Cc*OMT3 | | | | |
| *Cc*OMT4 | | | | |
| *Cj*4′OMT | *Coptis japonica* | | | |
| *Cm*OMT1 | *Chelidonium majus* | | | |
| *Cm*OMT2 | | | | |
| *Cm*OMT1 | | | | |
| *Cs*OMT1 | *Corydalis saxicola* | | | |
| *Ec*OMT1 | *Eschscholzia californica* | | + | |
| *Ec*OMT2 (7OMT/SOMT) | | | + | |
| *Ec*OMT3 | | | | |
| *Ec*OMT4 | | | | |
| *Ec*OMT5 | | | + | |
| *Ec*OMT6 | | | | |
| *Gf*OMT1 | *Glaucium flavum* | | | |
| *Gf*OMT2 | | | | |
| *Gf*OMT3 | | | | |
| *Hc*OMT1 | *Hypericum calycinum* | | | |
| *Nd*OMT1 | *Nandina domestica* | | | |
| *Nn*6OMT1 | *Nelumbo nucifera* | | | |
| *Nn*6OMT2 | | | | |
| *Nn*6OMT3 | | | | |
| *Nn*6OMT4 | | | | |
| *Nn*7OMT1 | | | | |
| *Nn*7OMT2 | | | | |
| *Nn*7OMT3 | | | | |
| *Nn*OMT5 | | | + | |
| *Ns*OMT1 | *Nigella sativa* | | | |
| *Ns*OMT2 | | | | |
| *Pb*OMT1 | *Papaver bracteatum* | | | |
| *Pb*OMT2 | | | | |
| *Pb*OMT3 | | | | |
| *Pb*OMT4 | | | | |
| *Ps*OMT1 (*Ps*N7OMT) | *Papaver somniferum* | | + | |

| | | | | |
|---|---|---|---|---|
| *Ps*OMT2 | | | | + |
| *Ps*OMT3 | | | | |
| *Ps*OMT4 | | | | |
| *Ps*OMT5 | | | | |
| *Ps*OMT6 | | | | |
| *Ps*OMT7 | | | | |
| *Ps*OMT8 | | | | |
| *Ps*OMT9 | | | | |
| *Ps*OMT10 | | | | |
| *Ps*OMT11 | | | | |
| *Ps*OMT12 | | | | |
| *Ps*OMT13 | | | | |
| *Ps*OMT14 | | | | + |
| *Ps*OMT15 | | | | |
| *Ps*OMT16 | | | + | |
| *Ps*OMT17 | | + | | |
| *Ps*OMT18 (*Ps*6OMT) | | + | | |
| *Ps*OMT19 (*Ps*4′OMT2) | | | | + |
| *Ps*OMT20 | | | | + |
| *Ps*OMT21 | | | | |
| *Sc*OMT1 | *Sanguinaria canadensis* | | | |
| *Sc*OMT2 | | | | |
| *Sd*OMT1 | *Stylophorum diphyllum* | | | |
| *Tc*OMT1 | *Tinospora cordifolia* | | | |
| *Tf*OMT1 | *Thalictrum flavum* | | | |
| *Tf*OMT2 | | | | |
| *Vha*OMT1 | | | | |
| *Xs*OMT1 | *Xanthoriza simplicissima* | | | |

**Supplementary Table 2 – Plasmids utilized in this study**

| Plasmid | Description | Source or reference |
|---|---|---|
| pBOT-HIS | CEN6/ARS4$^{ori}$, pMB1$^{ori}$, Amp$^{R}$ , Kan$^{R}$ , *HIS3*, P$_{TEF1}$-GFP-T$_{PGI1}$ | [1] |
| pCAS-G418 | P$_{RNR2}$-*cas9*$_{NLS}$-T$_{CYC1}$, pUC, 2μ, P$_{tRNA_Tyr}$-3′HDV-gRNA-Scaffold-T$_{SNR52}$, P$_{TEF1}$-*kanMX*-T$_{TEF1}$ | [2] |
| pCAS-Hyg | P$_{RNR2}$-*cas9*$_{NLS}$-T$_{CYC1}$, pUC, 2μ, P$_{tRNA_Tyr}$-3′HDV-gRNA-Scaffold-T$_{SNR52}$, P$_{TEF1}$-*HphNTI*-T$_{TEF1}$ | [3] |
| pPSG450 | CEN6/ARS4$^{ori}$, ColE1, Kan$^{R}$, *HIS3*, P$_{TEF1}$-*EcNMCH*-T$_{TDH3}$, P$_{TDH3}$-*Ps6OMT*-T$_{ADH1}$, P$_{PGK1}$-*Ps4'OMT2*-T$_{ENO1}$, P$_{TEF2}$-*PsCNMT*-T$_{SSA1}$, P$_{HHF1}$-*AtATR2*-T$_{ENO2}$, | [3] |
| pHUM | CEN6/ARS4$^{ori}$, ColE1, Amp$^{R}$, *HIS3*, *URA3*, *MET17* | [4] |
| pHUM-PrDRR | pHUM backbone, P$_{TDH3}$-*Pr*DRR-T$_{EFM1}$ | This study |
| pHUM-PsDRR | pHUM backbone, P$_{TDH3}$-*Ps*DRR-T$_{EFM1}$ | This study |
| pHUM-PbDRR1 | pHUM backbone, P$_{TDH3}$-*Pb*DRR1-T$_{EFM1}$ | This study |
| pHUM-PbDRR2 | pHUM backbone, P$_{TDH3}$-*Pb*DRR2-T$_{EFM1}$ | This study |
| pHUM-PrDRS-PsDRR | pHUM backbone, P$_{TDH3}$-*Pr*DRS-T$_{IDP1}$, P$_{CCW12}$-*Ps*DRR-T$_{EFM1}$ | This study |

**Supplementary Table 3 – Strains utilized in this study**

| ID | Description | Parent | Locus+manipulation | Reference |
|---|---|---|---|---|
| LP478 | (*S*)-Norcoclaurine strain | NA | NA | [3] |
| LP603 | (*S*)-Norcoclaurine strain | LP478 | *gre2*Δ | This study |
| LB604 | (*S*)-Coclaurine strain | LB603 | USERX-3-*Ps6OMT*-USERX-3 | This study |
| LP605 | (*S*)-*N*-Methylcoclaurine strain | LP603 | USERX-3-*Ps6OMT-PsCNMT*-USERX-3 | This study |
| LP609 | (*S*)-Armepavine strain | LP605 | USERXI-2-*NnOMT5*-USERXI-2 | This study |
| LP624 | *De novo* A9 pronuciferine assembly | LP609 | USERX-1-*NnCYP80G-AtATR2-AtCYB5D*-USERX-1 | This study |
| LP626 | BY4741 C1 pronuciferine assembly | BY4741 | USERX-1-*NnCYP80G-AtATR2-CrCYB5*-USERX-1 | This study |
| LP627 | Deletion of *NnOMT5* from BY4741 pronuciferine assembly | LP626 | Δ*NnOMT5* | This study |
| LP625 | *De novo* G5 pronuciferine assembly | LP609 | USERX-1-*NnCYP80A-AtATR2-PbCYB5*-USERX-1 | This study |
| LP615 | BY4741 3× MTase strain | BY4741 | USERX-3-*Ps6OMT-PsCNMT*-USERX-3<br>USERXI-2-*NnOMT5*-USERXI-2 | This study |
| LP623 | *De novo* A2 pronuciferine assembly | LP609 | USERX-1-*LsCYP80A-CcCPR2-AtCYB5D*-USERX-1 | This study |
| LP634 | Chromosomal *PrDRS-PsDRR* in A2 pronuciferine assembly | LP623 | 416d-*PrDRS-PsDRR*-416d | This study |
| LP631 | Deletion of *NnOMT5* from G5 *Nn*CYP80A assembly | LP625 | Δ*NnOMT5* | This study |
| LP636 | Chromosomal *PrDRS-PsDRR* in G5 *Nn*CYP80A Δ*NnOMT5* assembly | LP631 | 416d-*PrDRS-PsDRR*-416d | This study |
| LP638 | Chromosomal *PrDRS-PsDRR* in (*S*)-*N*-Methylcoclaurine strain | LP605 | 416d-*PrDRS-PsDRR*-416d | This study |
| LP637 | *Bs*CYP80A1+*Hc*CPR host | LP638 | USERXI-2-*BsCYP80A1-HcCPR*-USERXI-2 | This study |

**Supplementary Table 4 – Expression cassettes utilized in this study**

| Description | Cassette | Locus |
|---|---|---|
| *Ps6OMT* + *PsCNMT* | $P_{TEF1}$-*Ps6OMT*-$T_{VMA2}$-LTP1-$P_{CCW12}$-*PsCNMT*-$T_{EFM1t}$ | USERX-3 |
| *NnOMT5* | $P_{TDH3}$-*NnOMT5*-$T_{TDH1}$ | USERXI-2 |
| *Nn6OMT1* | $P_{TDH3}$-*Nn6OMT1*-$T_{TDH1}$ | 106a |
| *Nn6OMT2* | $P_{TDH3}$-*Nn6OMT2*-$T_{TDH1}$ | 106a |
| *Nn6OMT3* | $P_{TDH3}$-*Nn6OMT3*-$T_{TDH1}$ | 106a |
| *Nn6OMT4* | $P_{TDH3}$-*Nn6OMT4*-$T_{TDH1}$ | 106a |
| *Ps6OMT* | $P_{TDH3}$-*Ps6OMT*-$T_{TDH1}$ | 106a |
| *NnCNMT1* | $P_{TDH3}$-*NnCNMT1*-$T_{TDH1}$ | 106a |
| *NnCNMT3* | $P_{TDH3}$-*NnCNMT3*-$T_{TDH1}$ | 106a |
| *PsCNMT* | $P_{TDH3}$-*PsCNMT*-$T_{TDH1}$ | 106a |
| *CmDRS* | $P_{TDH3}$-*CmDRS*-$T_{IDP1}$ | FgF 21 |
| *PbDRS1* | $P_{TDH3}$-*PbDRS1*-$T_{IDP1}$ | FgF 21 |
| *PbDRS2* | $P_{TDH3}$-*PbDRS2*-$T_{IDP1}$ | FgF 21 |
| *PbDRS3* | $P_{TDH3}$-*PbDRS3*-$T_{IDP1}$ | FgF 21 |
| *PbDRS4* | $P_{TDH3}$-*PbDRS4*-$T_{IDP1}$ | FgF 21 |
| *PrDRS* | $P_{TDH3}$-*PrDRS*-$T_{IDP1}$ | FgF 21 |
| *PsDRS* | $P_{TDH3}$-*PsDRS*-$T_{IDP1}$ | FgF 21 |
| *PbDRR1* | $P_{TDH3}$-*PbDRR1*-$T_{EFM1}$ | pHUM plasmid |
| *PbDRR2* | $P_{TDH3}$-*PbDRR2*-$T_{EFM1}$ | pHUM plasmid |
| *PrDRR* | $P_{TDH3}$-*PrDRR*-$T_{EFM1}$ | pHUM plasmid |
| *PsDRR* | $P_{TDH3}$-*PsDRR*-$T_{EFM1}$ | pHUM plasmid |
| *PrDRS-PsDRR* | $P_{TDH3}$-*PrDRS*-$T_{IDP1}$-LTP1-$P_{CCW12}$-*PsDRR*-$T_{EFM1}$ | 416d |
| *NnCYP80A* | $P_{TDH3}$-*NnCYP80A*-$T_{ATG10}$ | USERX-1 |
| *NnCYP80G* | $P_{TDH3}$-*NnCYP80G*-$T_{ATG10}$ | USERX-1 |
| *LsCYP80G* | $P_{TDH3}$-*LsCYP80G*-$T_{ATG10}$ | USERX-1 |
| *AtATR2* | $P_{TDH3}$-*AtATR2*-$T_{ATG10}$ | USERX-1 |
| *CcCPR2* | $P_{TDH3}$-*CcCPR2*-$T_{ATG10}$ | USERX-1 |
| *HcCPR* | $P_{TDH3}$-*HcCPR*-$T_{ATG10}$ | USERX-1 |
| *PsCPR* | $P_{TDH3}$-*PsCPR*-$T_{ATG10}$ | USERX-1 |
| *AtCYB5D* | $P_{TDH3}$-*AtCYB5D*-$T_{ATG10}$ | USERX-1 |
| *CrCYB5* | $P_{TDH3}$-*CrCYB5*-$T_{ATG10}$ | USERX-1 |
| *EcCYB5* | $P_{TDH3}$-*EcCYB5*-$T_{ATG10}$ | USERX-1 |
| *PbCYB5* | $P_{TDH3}$-*PbCYB5*-$T_{ATG10}$ | USERX-1 |
| *BsCYP80A1* + *HcCPR* | $P_{TDH3}$-*BsCYP80A1*-$T_{ATG10}$-LTP1-$P_{HHF1}$-*HcCPR*-$T_{PGK1}$ | USERXI-2 |

**Supplementary Table 5 – *S. cerevisiae* integration sites utilized in this study**

| Target site ID | Target site sequence[a] | Reference |
|---|---|---|
| *GRE2* | TAGAATACGGAATTTTCTCGCGG | [3] |
| *NnOMT5* | TAACAGCTTTGCAGTTTTCGTGG | This study |
| USERX-1 | GTAGCTACAAGAACATATGGTGG | [5] |
| USERX-3 | GATCGCCGAATGGCACGCGAGGG | [5] |
| USERXI-2 | AGTTTAGATGTAGGTTTTAGCGG | [5] |
| USERXII-1 | CTTTGATTTTGGCATCGGTTCGG | [5] |
| 416d | TAGTGCACTTACCCCACGTTCGG | [6] |
| 511b | GGGTTTGGCACAATTTGGCTTGG | [6] |
| 911b | GTAATATTGTCTTGTTTCCCTGG | [6] |
| 106a | ATACGGTCAGGGTAGCGCCCTGG | [6] |

[a] PAMs are underlined.
